# The architecture, assembly, and evolution of a complex flagellar motor

**DOI:** 10.1101/2025.02.19.638559

**Authors:** Xueyin Feng, Shoichi Tachiyama, Jing He, Siqi Zhu, Hang Zhao, Jack M. Botting, Yanran Liu, Yuanyuan Chen, Canfeng Hua, María Lara-Tejero, Matthew A. B. Baker, Xiang Gao, Jun Liu, Beile Gao

## Abstract

Bacterial flagella drive motility in many species, likely including the last bacterial common ancestor ^1,2^. Knowledge of flagellar assembly and function has mainly come from studies of *Escherichia coli* and *Salmonella enterica*, which have simple flagellar motors ^3–7^. However, most flagellated bacteria possess complex motors with unique, species-specific adaptations whose mechanisms and evolution remain largely unexplored ^8–10^. Here, we deploy a multidisciplinary approach to build a near-complete model of the flagellar motor in *Campylobacter jejuni*, revealing its remarkable complexity in architecture and composition. We identify an E-ring around the MS-ring, a periplasmic cage with two distinctive conformations, and an intricate interaction network between the E-ring and cage. These scaffolds play critical roles in stabilizing and regulating 17 torque-generating stator complexes for optimal motility. In-depth evolutionary analyses uncover the ancient origin and prevalence of the E-ring in flagellated species of the domain *Bacteria* as well as a unique exaptation of type IV pili components PilMNOPQF in the ancestral motor of the phylum *Campylobacterota*. Collectively, our studies reveal novel mechanisms of assembly and function in complex flagellar motors and shed light on the evolution of flagella and modern bacterial species.

## Main text

Flagellar motors are complex nanomachines that are highly diverse across bacterial species ^8,11,12^. Among these, the classical models *E. coli* and *S. enterica* have the same, simple flagellar motor structures ^8,13^. Between the inner membrane and outer membrane of the *E. coli* model, the motor is composed of an MS-ring and central rod, which is surrounded by the P-ring and L-ring at the peptidoglycan layer and outer membrane, respectively ^14^. Motor rotation is driven by stator complexes that harness the proton motive force to generate torque ^15^. Stator complexes are highly dynamic in *E. coli* and *S. enterica* ^16^. Importantly, the flagellar motors in species of *Enterobacteriaceae*, including *E. coli* and *S. enterica*, were not inherited from the common ancestor of γ-proteobacteria but rather acquired by horizontal gene transfer from an ancestral β-proteobacterium ^17^.

By contrast, the motors of many flagellated species have auxiliary structures, in addition to the P-ring and L-ring, around the rod in the periplasm ^18–20^. Such motors with additional periplasmic scaffolds are termed complex motors and classified into three categories ^21^: (1) outer-membrane-associated scaffold motors, as in *Vibrio* spp. that have H/T/O-rings connected to the outer membrane ^20,22^, (2) inner-membrane-associated scaffold motors present in *Spirochaetota* species such as *Borrelia burgdorferi* (these scaffolds are known as “collars”) ^23–25^, and (3) integrated scaffold motors spanning the periplasm and associated with both the outer membrane and inner membrane, as in *C. jejuni* and *Helicobacter pylori* ^9,26,27^. Unlike the *E. coli* motor with its dynamic stator complexes ^16^, complex motors possess remarkable structural intricacy that appears to stabilize stator complexes in 18 out of 24 species (Supplementary Table 1). However, the evolution and function of complex motors remain largely unknown ^9,28,29^.

It was recently proposed that complex motors have evolved from a simple ancestral motor, whose prototype is the *E. coli* motor ^13,30^. However, this proposal was based on motor structures in 7∼8 species and on the assumption that the ancestral motor should be the simplest ^13,30^. Evidence from bacterial phylogeny does not align with this notion ^31^ and is confounded by differences in specific proteins and structures across species. To better understand the evolutionary path and mechanism of complex motors, the first challenge is to identify components that constitute the additional structures. The motility of the human pathogen *C. jejuni* is driven by a complex flagellar motor in each cell pole ^32^. The complex flagellar motor of *C. jejuni* has a well-studied “parts list”, thanks to multiple high-throughput transposon library screenings and extensive characterization of novel flagellar genes ^9,26,33–36^. Notably, our previous Tn-seq screenings using cell invasion and mouse infection identified several novel genes impacting motility, including those that, when deleted, result in a significant decrease in fitness during host interaction but no change in motility on artificial soft agar or liquid medium ^33,37^.

The *C. jejuni* motor contains three disks: a basal disk composed of FlgP just below the outer membrane ^9,38^, a medial disk composed of PflC and PflD ^26^, and a proximal disk containing PflA and PflB ^9,26^. These findings provided a foundation to dissect complex motors with homologs and suggested that complex motors possess mechanisms different from those in the classical model ^30,38,39^.

### Mapping the components in motor periplasmic scaffolds by imaging specific mutants

We structurally characterized new periplasmic scaffold proteins in the *C. jejuni* motor by using cryo-electron tomography (cryo-ET) and molecular genetics. Among multiple flagellar genes identified from Tn-seq screenings ^33,37^, mutants of *flgY* (*CJJ81176_1488* in *C. jejuni* 81-176 genome) ^33^ and three genes in a cluster (*CJJ81176_0481, 0480, 0479*) ^37^ showed loss of specific densities in the periplasmic region of the flagellar motor structures, with all other scaffolds remaining largely intact, as in wild type (Fig. 1a-d).

**Fig. 1.**
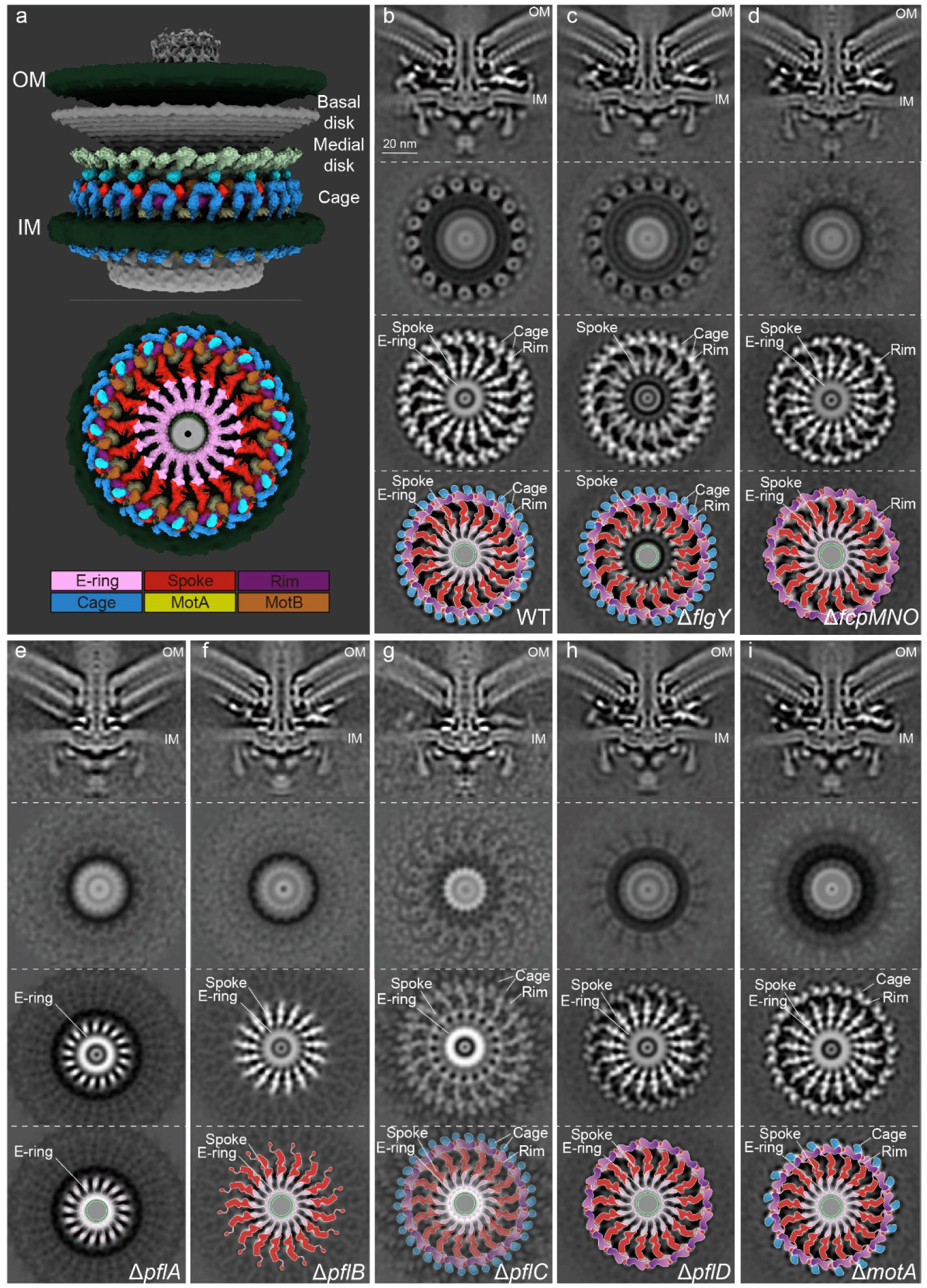
Mapping components in *C. jejuni* motor. **a**, Cut-through view of the organization of the *C. jejuni* motor in situ, with a focus on periplasmic scaffolds below the medial disk. Color scheme for proteins or complexes is indicated below and the E-ring in both light and dark pink represents dimer of FlgY. **b-i**, Each column with four images from top to bottom contains central section of motor structure, cross-section of stator region, cross-section of proximal disk region (refined), and cartoon representation of proximal disk region, from wild type or seven flagellar gene mutants. Scale bar in **b**, 20 nm, applies to columns **b** through **i**. Color scheme in cartoon representations follows the same as in **a**.

Compared to the wild-type motor, the Δ*flgY* motor lacks a periplasmic spoke-ring around the MS-ring (Fig. 1a-c), a location like that of the “E-ring” first discovered by electron microscopy in *Caulobacter crescentus* ^40,41^. Extensive classification and refinement of the periplasmic region of the wild-type *C. jejuni* motor structure revealed that the E-ring is not a continuous ring or disk. Instead, it is composed of a small, thin ring around the MS-ring and 17 separate spokes radiating from the small ring (Fig. 1b). These spokes connect in a 1:1 ratio with 17 longer, distal spokes that extend to 17 stator complexes and attach to a continuous rim-like structure (Fig. 1b). The distal spokes and rim are composed of PflA (spokes) and PflB (rim) ^26^. In addition, an N-terminal 17-aa signal peptide was predicted for FlgY (Extended Data Fig. 1a). Thus, FlgY is likely a periplasmic component of the E-ring.

The triple knockout mutant Δ*CJJ81176_0481-0479* lacks the peripheral structure most distal from the central rod in wild-type *C. jejuni* (Fig. 1a,b,d). This peripheral structure is made of 34 units embedded in the inner membrane and below the medial disk (Fig. 1a,b). The position and shape of this structure are very similar to those of the lower cage in *H. pylori*, which consists of PilM/PilN/PilO homologs of type IV pili (T4P) ^42^. Sequence analyses and AlphaFold3 structure prediction of CJJ81176_0481, 0480, and 0479 revealed that these three proteins in *C. jejuni* are homologs of *H. pylori* PilM/PilN/PilO (Extended Data Fig. 1b,c). Coimmunoprecipitation (co-IP) followed by liquid chromatography-mass spectrometry (LC-MS/MS) analysis also revealed that each protein can pull down the other one or two proteins in *C. jejuni*, suggesting that they form a complex (Supplementary Table 2). In addition, *C. jejuni* 81-176 does not encode other components of T4P, and these three genes show co-occurrence with the ancient flagellar gene set and F3 chemosensory class in genomes of *Campylobacterota* species (discussed in detail in later section). Hence, we named the proteins CJJ81176_0481, 0480, and 0479 **f**lagellar **<u>c</u>**age **<u>p</u>**roteins FcpM/FcpN/FcpO, based on their shared homology with PilM/PilN/PilO but different roles as flagellar components.

To explore the structural dependency of these new scaffolds with surrounding structures, such as the proximal and medial disks made of PflA/PflB/PflC/PflD, we compared the motor structures of four mutants: Δ*pflA*, Δ*pflB*, Δ*pflC*, and Δ*pflD* (Fig. 1e-h). In Δ*pflA*, only the E-ring is present with more plasticity in shape in the inner-membrane proximal region, while the PflA spokes, PflB rim, and cage units are absent (Fig. 1e). For Δ*pflB*, both the E-ring and spokes remain, but with less resolved densities, while the rim and cage units are missing (Fig. 1f). The Δ*pflD* mutant mimics Δ*fcpMNO*, missing the entire peripheral cage. (Fig. 1d,h). Though not as stable as those in the wild-type motor, the E-ring, spokes, and rim remain in the Δ*pflC* mutant (Fig. 1g), in contrast to the complete scaffold loss recently reported ^26^. Together, these findings establish that assembly of the E- ring is not dependent on PflA/PflB/PflC/PflD or FcpM/FcpN/FcpO and that the full set of 34 peripheral cage units requires all scaffolds inside (PflA/PflB/PflC/PflD) except the E- ring.

To determine the roles of the scaffolds in motor function and bacterial motility, we compared the stator densities of seven mutants and wild-type *C. jejuni* (Fig.1b-i). The Δ*flgY* mutant showed the same stator density as wild type, with approximately 80% of the motors having a full stator ring and remaining 20% having less-resolved stator densities (Fig. 1b,c and Extended Data Fig. 2a). By contrast, neither the Δ*pflA* nor Δ*pflB* mutant has stator densities in the motor structure, similar to Δ*motA* (Fig. 1e,f,i). The three mutants Δ*pflC,* Δ*pflD*, and Δ*fcpMNO* have reduced stator densities (Fig. 1d,g,h). Specifically, approximately 10% of the Δ*pflD* and Δ*fcpMNO* mutants have stator complexes loaded in the motor, and the Δ*pflC* mutant has least stator complexes in the motor (Extended Data Fig .2a and Fig. 1d,g,h). Importantly, the differences in stator density/occupancy among these mutants are consistent with their motility phenotypes on soft agar, including the greatly reduced but not abolished motility of the Δ*pflC* mutant (Extended Data Fig. 2b). Collectively, our data provide direct evidence that the inner-membrane scaffolds are critical for stator assembly and motility in *C. jejuni* (Fig. 1 and Extended Data Fig. 2).

### FlgY dimers form the E-ring around the MS-ring via ARM-like domains

Next, we dissected the structure, function, and interaction of the inner-membrane proximal scaffolds at the protein or domain/motif level. Starting from the innermost E-ring, candidate component FlgY was structurally analyzed. AlphaFold3 prediction showed that FlgY is entirely α-helical, with the N-terminal 15-116 aa forming a long α-helix and the remaining 56 residues comprising a four-membered, right-handed superhelix (Fig. 2a). The superhelical domain contains a hydrophobic core, showing structural similarity to the N-terminal cytosolic domain of the Mg^2+^ transporter MgtE as well as to the armadillo repeat motif (ARM)-like motifs in flagellar rotor protein FliG (Extended Data Fig. 3a) ^43–45^. In particular, this C-terminal domain, here named FlgY_ARM_, differs from the canonical ARM repeat with regard to helical packing ^44,45^ and matches well with the ARM-like motif of MgtE and FliG (Extended Data Fig. 3a,b).

**Fig. 2.**
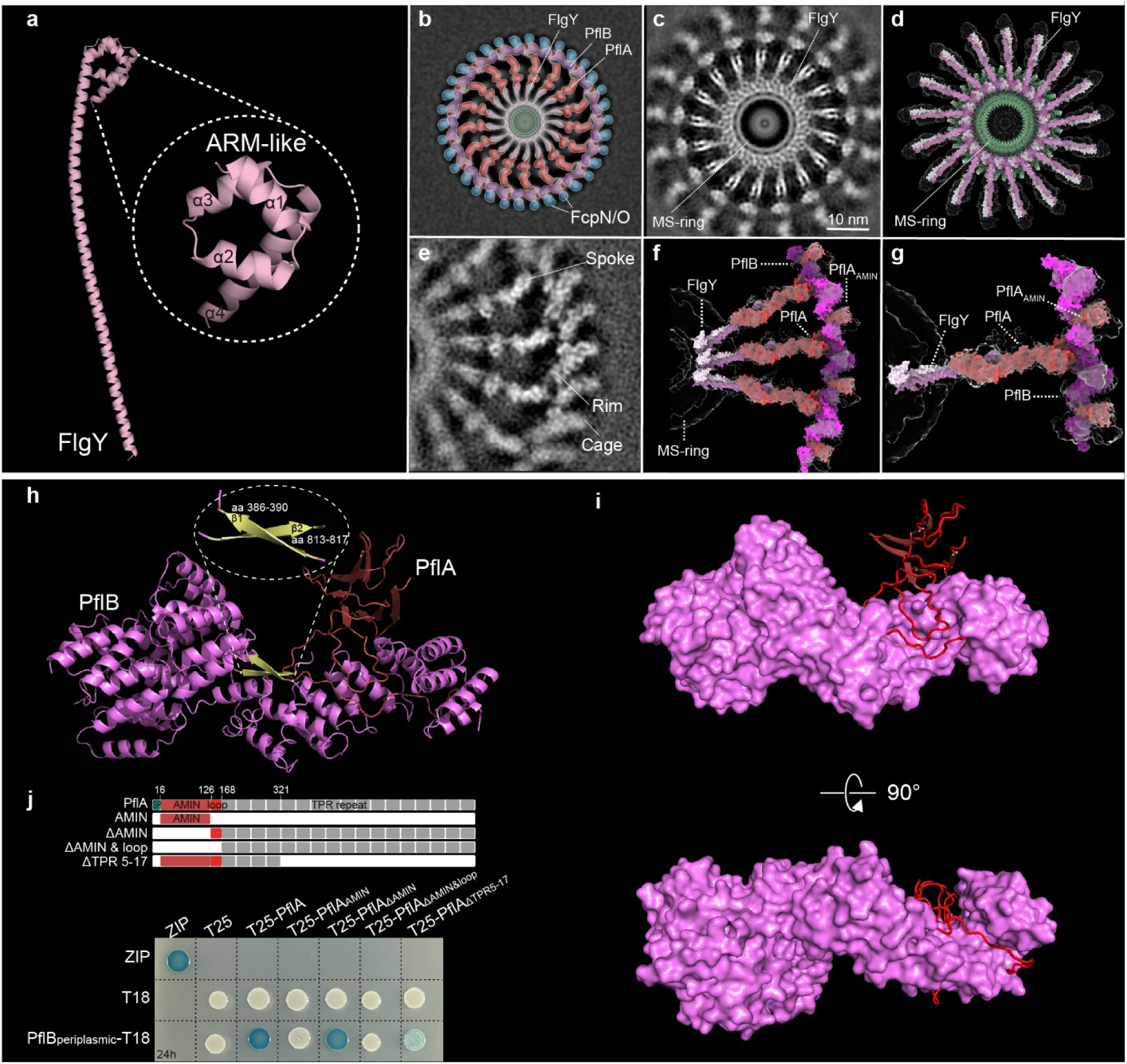
Dissecting the E-ring, spokes and rim. **a**, AlphaFold3-predicted structure of FlgY, with an inset showing the C-terminal ARM-like domain. **b,** Cartoon model of cross-section of motor structure from wild type, showing E-ring, spokes, rim, and cage units. Color scheme follows the same as in **Fig.1a**. **c**, Local refinement of periplasmic region in panel **b** revealed high-resolution structure of E-ring. **d**, Top view of E- ring model with C17 symmetry produced by docking predicted structure of FlgY dimer (light/dark pink, more details see Extended Data Fig. 3e) into cryo-ET density (transparent regions). **e**, Local refinement of periplasmic region in panel **b** revealed high-resolution structure of PflA spokes, PflB rim and cage. **f**, Modelled FlgY dimer and PflAB complex (red and magenta) docked into refined map **(e**). **g**, Close-up view of single docked unit shown in **f,** highlighting PflA_AMIN_ domain at the rim region. **h**, Cartoon representation of the single-particle cryo-EM structure of PflA (red) and PflB (magenta and yellow for β-sheet) complex. **i**, Cartoon representation of PflA structure (red) and surface presentation of PflB structure (magenta) to show how the PflA loop wraps around PflB. **j,** BTH analysis to dissect the interaction mode of PflA and PflB at domain or motif level. The domain organization of full-length PflA and various truncation variants of PflA made for BTH analyses were indicated on the top.

Purified FlgY without the predicted signal peptide (FlgY_15-172_) forms a dimer in solution (Extended Data Fig. 3c). In addition, variants with different truncations in the N-terminal long α-helix region still form a dimer, suggesting that the FlgY dimer is stable even without the long α-helix region (Extended Data Fig. 3c). ARM repeats stack on each other intramolecularly or intermolecularly, as shown for ARM-like motifs in FliG that drive C-ring formation ^46,47^. To test whether FlgY_ARM_ shares the same feature, crosslinking experiments were carried out on FlgY_15-172_ and FlgY_ARM_. Both can form higher oligomers (Extended Data Fig. 3d), suggesting that dimeric FlgY_ARM_ may stack with an adjacent FlgY_ARM_ as well. We then docked the predicted structure of the FlgY dimer into a refined cryo-ET map at 12 Å resolution (Fig. 2b-d and Extended Data Fig. 3e). 34 ARM-like domains from 17 FlgY dimers form a ring around the β-collar of the MS-ring, and each dimeric coiled coil domain points outward, interacting with the distal spoke formed by PflA (Fig. 2d-g). This interaction between FlgY and PflA is consistent with the additional densities at the inner end of the PflA spokes in the Δ*flgY* mutant (Fig. 2f,g and 1c). Additionally, the interaction of PflA with FlgY_15-172_, but not with FlgY_ARM_ without the N-terminal long α-helix, was detected by microscale thermophoresis (Extended Data Fig. 3f). These results suggest that FlgY dimers constitute the E-ring and connect with PflA spokes via the long α-helix region.

### PflA spokes tightly bind the PflB rim

To determine how the spokes interact with the rim, we improved *in-situ* structure of the spoke-rim complex at 12 Å resolution by 17-fold symmetry expansion and focused refinement (Fig. 2e) in addition to solving a single particle cryo-EM structure of the PflAB complex at 3.23 Å resolution (Fig. 2h,i, Extended Data Fig. 4). A combination of both *in-situ* and *in-vitro* structures with AlphaFold3 prediction of PflA enabled us to build the pseudoatomic model of the spoke-rim complex (Fig. 2f,g). In this model, PflA and PflB are in a 1:1 ratio, with the majority of PflB (178-820 aa) resolved at near-atomic resolution (Fig. 2h). PflB is mainly composed of tetratricopeptide (TPR) repeats and can be divided into two parts by a β1-sheet (aa 386-390) (Fig. 2h). The N-terminal TPR repeats before the β1-sheet are relatively extended, while those after the β1-sheet are tightly packed (Fig. 2h). Notably, the last 5 C-terminal residues form a β2-sheet (aa 813-817) that makes an antiparallel β-pair with the β1-sheet (Fig. 2h). This antiparallel β-pair resembles a “hook and loop”, as the two β-sheets are separated by 423 residues in sequence yet engaged together in the 3D structure (Fig. 2h). We hypothesize that this β-pair stabilizes the TPR repeats after the β1-sheet, making it less extended than the N-terminus in overall shape (Fig. 2h).

PflA consists of the N-terminal β-sandwich domain, flexible loop, and C-terminal 17 repetitive TPR repeats. The N-terminal β-sandwich domain and flexible loop region (16- 163 aa) are resolved in our cryo-EM structure (Fig. 2h,i). The β-sandwich domain of PflA is made of four pairs of antiparallel β-sheets and here named PflA_AMIN_ for its resemblance to the AMIN1 and AMIN2 domains of PilQ in T4P (Extended Data Fig. 5a). Interestingly, the long loop of PflA (126-163 aa) wraps around the N-terminal extended TPR repeats of PflB, mainly via hydrophobic contacts (Fig. 2i and Extended Data Fig. 5b). The critical role of this PflA loop in PflB interaction was further confirmed by bacterial two hybrid (BTH) analyses (Fig. 2j). Overall, the PflAB complex is akin to a “wire-wrapped pendant”, with PflA (wire) looping around PflB (stone), forming a tight interaction (Fig. 2i) and assembling the PflAB spoke-rim (Fig. 2g).

To dissect the role of each domain and/or motif in the PflAB complex, various truncations of PflA or PflB were made and complemented into the Δ*pflA* or Δ*pflB* mutant, respectively, for soft agar motility assays (Extended Data Fig. 5c). Results show that PflA_AMIN_ and the first 15 TPR repeats are important for PflA function because their deletion results in a non-motile phenotype (Extended Data Fig. 5c). Our PflAB complex structure suggests that these two regions are not involved in PflB interaction, a finding further confirmed by BTH analyses (Fig. 2h-j). Notably, PflA was detected in the co-IP product of 3xFLAG-tagged FcpO (Supplementary Table 2), implying that PflA spokes interact with the cage units in addition to the rim. To test this notion, we performed BTH analyses for PflA and the PflA_AMIN_ domain with multiple scaffolding proteins in the peripheral region. Results show that the PflA_AMIN_ domain, but not the whole periplasmic region of PflA, can interact with FcpO and FliL (Extended Data Fig. 5d).

Finally, removal of the N-terminal cytoplasmic region of PflB reduced motility to 50% of wild-type levels on soft agar, suggesting that this region plays a role in PflB function, perhaps interacting with other protein(s) in the cytosol (Extended Data Fig. 5c). Deletion of the β2-sheet from PflB results in complete loss of motility (Extended Data Fig. 5c), supporting a structural role for the β-pair in PflB (Fig. 2h).

### Two PflD conformations form different tetrameric cage units with FcpMNO

Our recent study revealed that the number of peripheral cage units is reduced to 17 in both the Δ*motA* and Δ*flgX* mutants, half the number in the wild-type motor (Fig. 1i) ^48^. We proposed that half of the cage units assemble after stator association, but it remained unknown how the FcpMNO proteins might achieve this two-step assembly. Here, we compared the refined structures of the peripheral cage region of wild type and three mutants: Δ*fcpMNO*, Δ*motA*, and Δ*pflD* (Fig. 3a). In wild type and the Δ*fcpMNO* and Δ*motA* mutants, 17 density spots are present below the medial disk (just below the PflC_2-6_ radial spokes ^26^) but missing from the Δ*pflD* mutant (Fig. 3a and Extended Data Fig. 6a). Therefore, these 17 spots likely represent PflD, consistent with the proposed position for PflD in a recent study that also indicated a density of unknown composition next to PflD ^26^. We examined this unknown density between two cage units and found that it is absent from all three mutants, including Δ*motA*, suggesting that it is related to the missing cage units and stator complexes in the Δ*motA* mutant (Fig. 3a and Extended Data Fig. 6a).

**Fig. 3.**
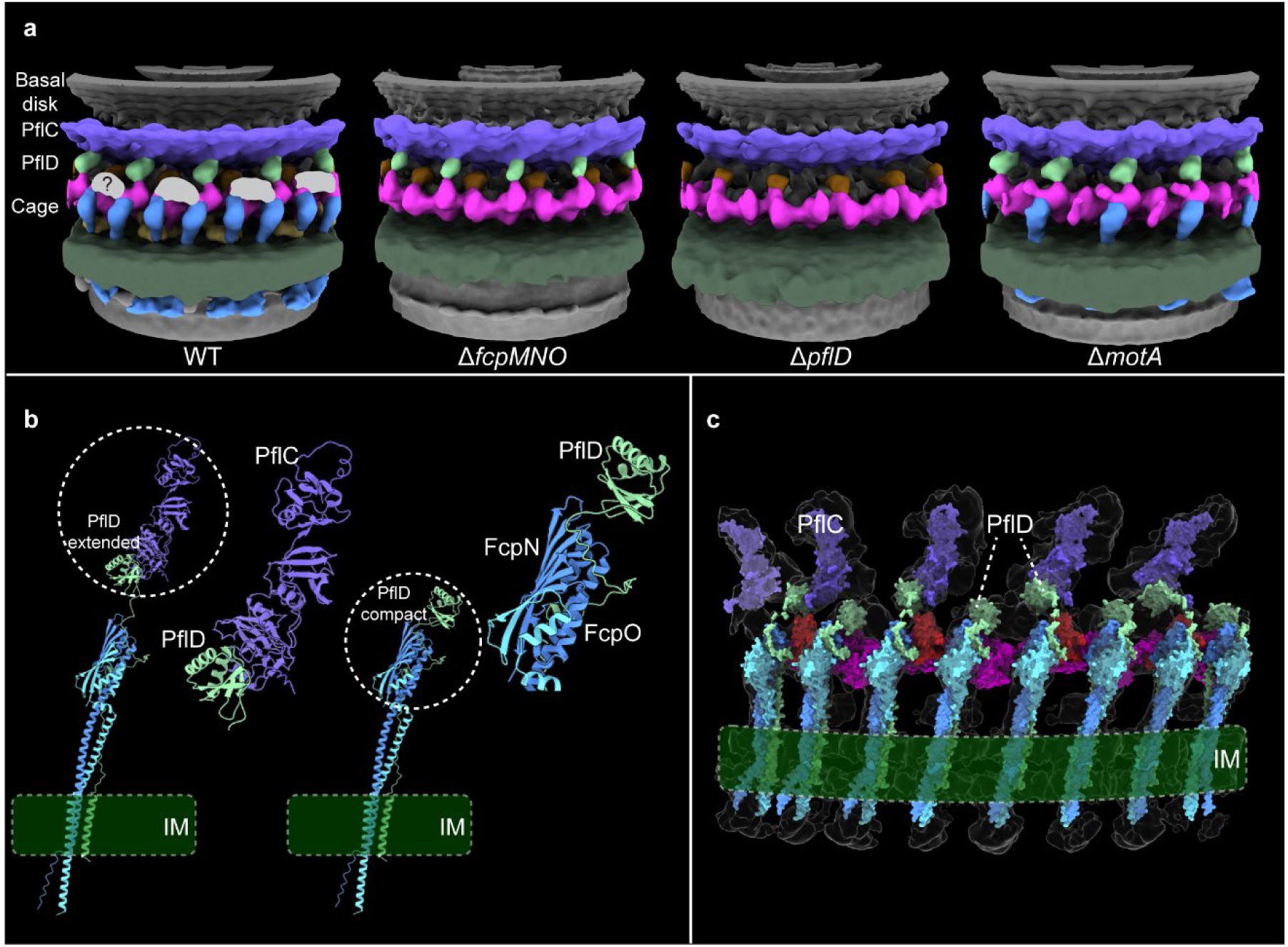
PflD in two conformations form different tetrameric cage units with FcpMNO. **a**, Comparison of 3D views of motor structures (isosurface rendering) from wild type and three mutants Δ*fcpMNO*, Δ*pflD*, and Δ*motA*. Below the medial disk (purple), PflD is highlighted in green, PflB rim in magenta, and cage units in blue. The unknown density between two cage units next to PflD position is in grey and marked with question mark (?). **b**, AlphaFold-Multimer predicted structures of the PflD (light green)-FcpNO (dark and light blue) complex with two conformations. Left: PflD with extended PflD_PilP-loop_ connecting PflC spoke (purple) by PflD_PilQ-N1_ domain; right: PflD with contracted PflD_PilP-loop_ making its PflD_PilQ-N1_ domain closer to FcpNO and detaching from PflC spoke. **c**, PflD-FcpNO complexes in two conformations fitted into the segmented cryo-ET map (transparent).

PflD contains an N-terminal transmembrane motif, and its structure predicted by AlphaFold3 has a C-terminal globular domain similar to the N1 domain of PilQ from T4P ^49^ (Extended Data Fig. 6b). Between the transmembrane motif and C-terminal PflD_PilQ-N1_ domain is a 60 aa flexible loop very similar to the loop region of PilP (named PflD_PilP-loop_) ^49^ (Extended Data Fig. 6b). A recent study on the T4P structure of *Pseudomonas aeruginosa* suggested that PilP interacts with PilNO via its short α-helix, together forming a tri-helix bundle-like structure ^49^. Alphafold3 prediction for PflD and FcpNO also showed a tri-helix bundle, but one formed by the N-terminal transmembrane segments of all three proteins (Extended Data Fig. 6c). This predicted complex appears more stable than PilNOP because PilP lacks a transmembrane motif and has a limited interaction interface with PilNO ^49^ (Extended Data Fig. 6c).

As a flexible loop, PflD_PilP-loop_ can be stretched or contracted, leading us to speculate that PflD in contracted form is the unknown component between the two cage units in wild type (Fig. 3b). To test this notion, the predicted PflD-FcpNO trimer structures in both stretched and contracted conformations were docked into our cryo-ET density map of the wild-type motor in alternating order (Fig. 3b,c). Both conformations fit very well with the periphery structure below the medial disk, specifically with the PflD_PilQ-N1_ domain positioned in the 17 density spots as recently proposed ^26^ and also the other 17 densities between two cage units (Fig. 3c). We also detected interaction between PflD and FcpN and PflD and FcpO with all their transmembrane motifs included using BTH analyses, but no interaction between these protein pairs with only their periplasmic regions (Extended Data Fig. 6d). Therefore, we propose that the peripheral cage units are heteromeric tetramers made of FcpMNO and PflD, with PflD in alternating extended and contracted conformations. Moreover, the cryo-ET structure suggests that PflD with extended PflD_PilP- loop_ reaches to the PflC_2-6_ spokes above via its PflD_PilQ-N1_ domain, possibly linking 17 of the cage units to the medial disk (Fig. 3b,c). This linkage may be one reason why the half of the cage units with extended PflD_PilP-loop_ remain in the motor upon deletion of *motA*, while the other half fail to assemble in the absence of stator complexes (Fig. 3a).

### The model and dynamics of the complex flagellar motor in *C. jejuni*

The above findings by the integrated approach allowed us to build a near-complete pseudoatomic structure of the *C. jejuni* motor (Fig. 4a). In this model, FlgY dimers, together with PflA monomers, form 17 spokes radiating from the MS-ring to connect 17 stator complexes in the motor (Fig. 4b,c). Around the stator complexes, one L-shaped PflA binds one PflB via its flexible loop, locking 17 PflB monomers into a continuous rim that holds 17 MotB dimers (Fig. 4b,c). Peripheral to the stator ring and rim, 34 cage units made of FcpMNO and PflD further enclose the stator complexes (Fig. 4b,c). Towards the top section, 17 cage units with extended PflD_PilP-loop_ connect with 17 PflC_2-6_ spokes in the medial disk (Fig. 4a). Towards the bottom section, all proteins PflB, FliL, FcpN, FcpO, and PflD have a transmembrane motif, and thus the scaffolds they form include one rim, 17 FliL rings, and 34 cage units all anchored in the inner membrane (Fig. 4a). In addition, these proteins together with PflA form a complex interaction network, as suggested by our BTH analyses (Extended Data Fig. 5d and 6d). Overall, this scaffolding platform resembles a lattice (such as the “Hanging Gardens of Babylon”) with load-bearing beams (FlgY/PflA) and multiple pillars (PflB/FliL/FcpNO/PflD) embedded in the inner membrane, stably accommodating 17 stator complexes within the *C. jejuni* motor (Fig. 4a,b).

**Fig. 4.**
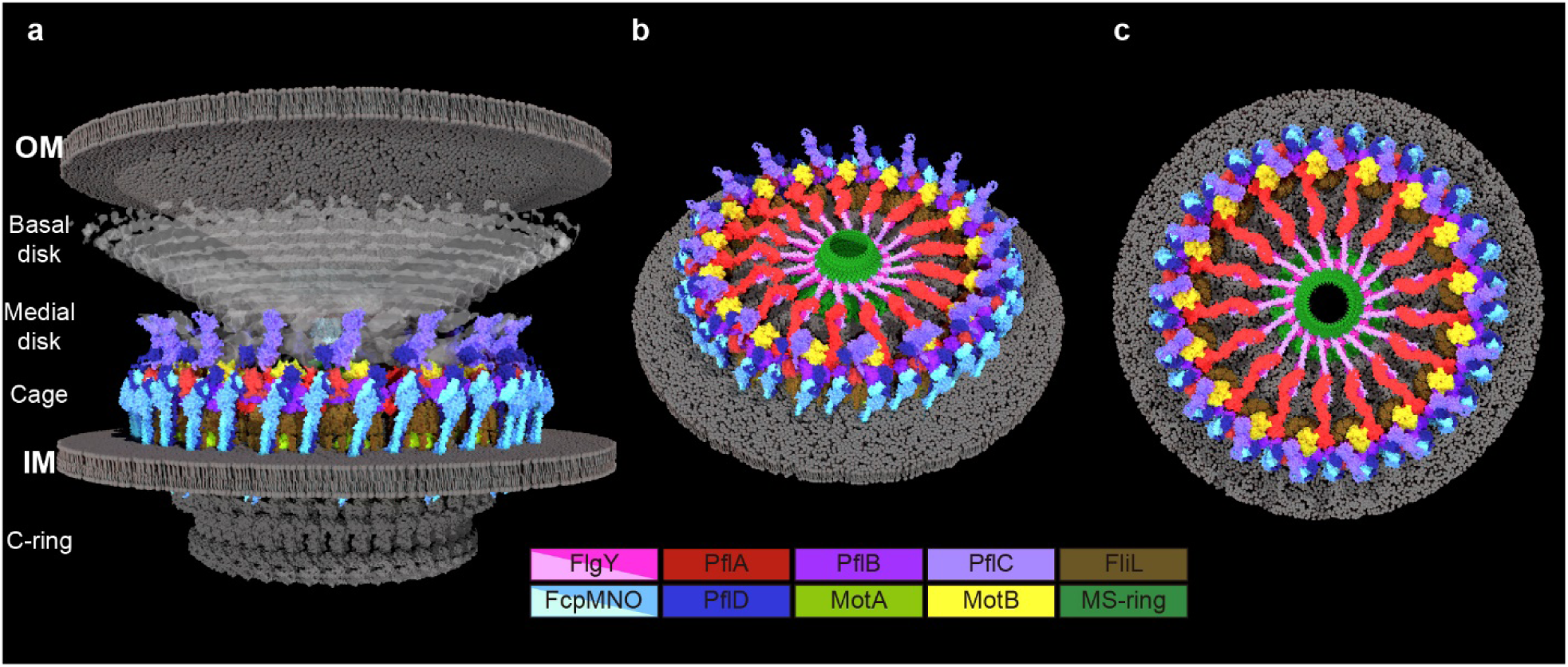
Model of *C. jejuni* flagellar motor. **a**, Cut-through view of *C. jejuni* motor built by docking structures of components listed below. Color scheme is also indicated below and structure source for proteins or complexes are indicated in Materials and Method. **b-c**, Tilted and top view of *C. jejuni* motor, removing top part to reveal the inner-membrane proximal scaffold components.

Notably, the peptidoglycan layer is not visible inside the motor. It seems clear that the periplasmic scaffolding platform encloses the stator complexes while excludes the peptidoglycan layer (Fig. 1b, 4a). Intriguingly, PflA contains a peptidoglycan-binding module, PflA_AMIN_ domain ^49–51^, which could remain in contact with the peptidoglycan layer through the gaps between the cage subunits (Fig. 4). By contrast, MotB is completely separated from the peptidoglycan layer due to the presence of the periplasmic scaffolds including FliL ring, PflA spokes, PflB rim, and FcpMNO/PflD cage (Fig. 4a). Therefore, the *C. jejuni* motor must utilize a different mechanism to control stator activation without directly interacting the peptidoglycan layer.

Importantly, focused classification enabled us to uncover conformational changes *en bloc* in wild-type *C. jejuni* motors (Supplementary Video 1, 2). Specifically, focused classification on the periplasmic scaffolds revealed their variable height relative to the MS-ring and rod, reflecting plasticity and adaptation of the motor to accommodate slight variation in the bacterial envelope (Extended Data Fig. 7a-d and Supplementary Video 1). Focused classification on the C-ring not only revealed its 40 subunits but captured their distinct orientations (Extended Data Fig. 7e and Supplementary Video 2), different from the 38-fold symmetry recently reported for *C. jejuni* C-ring ^26^. Given that the wild-type motors rotate constantly, a sequentially arranged image stack enabled us to visualize the motor rotation and dynamic fluctuations for the first time (Supplementary Video 2).

### Inner-membrane proximal scaffolds and stator complexes assemble before the rod in *C. jejuni*

In the classical model, assembly follows an inside-out sequence across the cell envelope ^52^. For complex motors, investigations of how and in what order the periplasmic scaffolds assemble are limited to a pilot study in *B. burgdorferi* ^53^. In *C. jejuni*, RpoN (σ^54^) and FliA (σ^28^) are specific regulators for flagellar gene expression ^32,54^. To explore the gene regulation of the inner-membrane proximal scaffolds, we performed RNA-seq on wild-type *C. jejuni* as well as Δ*rpoN* and Δ*fliA* mutants. Compared to the wild-type transcriptome, the expression levels of *flgY, pflA, pflB*, and *fcpM* remained the same in both mutants, while *fcpN*, *fcpO* and *pflD* showed a 2-fold decrease in the Δ*rpoN* mutant (Fig. 5a and Supplementary Table 3). The relative transcript abundance of *fcpN* and *fcpO* was very low in all samples examined (Supplementary Table 3), so we performed qPCR analyses for all seven genes to verify the RNA-seq data (Extended Data Fig. 8a). Our qPCR results show that none of these genes encoding for inner-membrane proximal scaffolds are regulated by RpoN or FliA (Extended Data Fig. 8a). In addition, no RpoN- or FliA-binding motifs were found in the promoter regions of *flgY*, *pflA*, *pflB*, *pflD*, or *fcpMNO* (Extended Data Fig. 8b). Therefore, genes encoding the E-ring, spokes, rim, and cage are outside the known flagellar transcriptional cascade in *C. jejuni* (Fig. 5b).

**Fig. 5.**
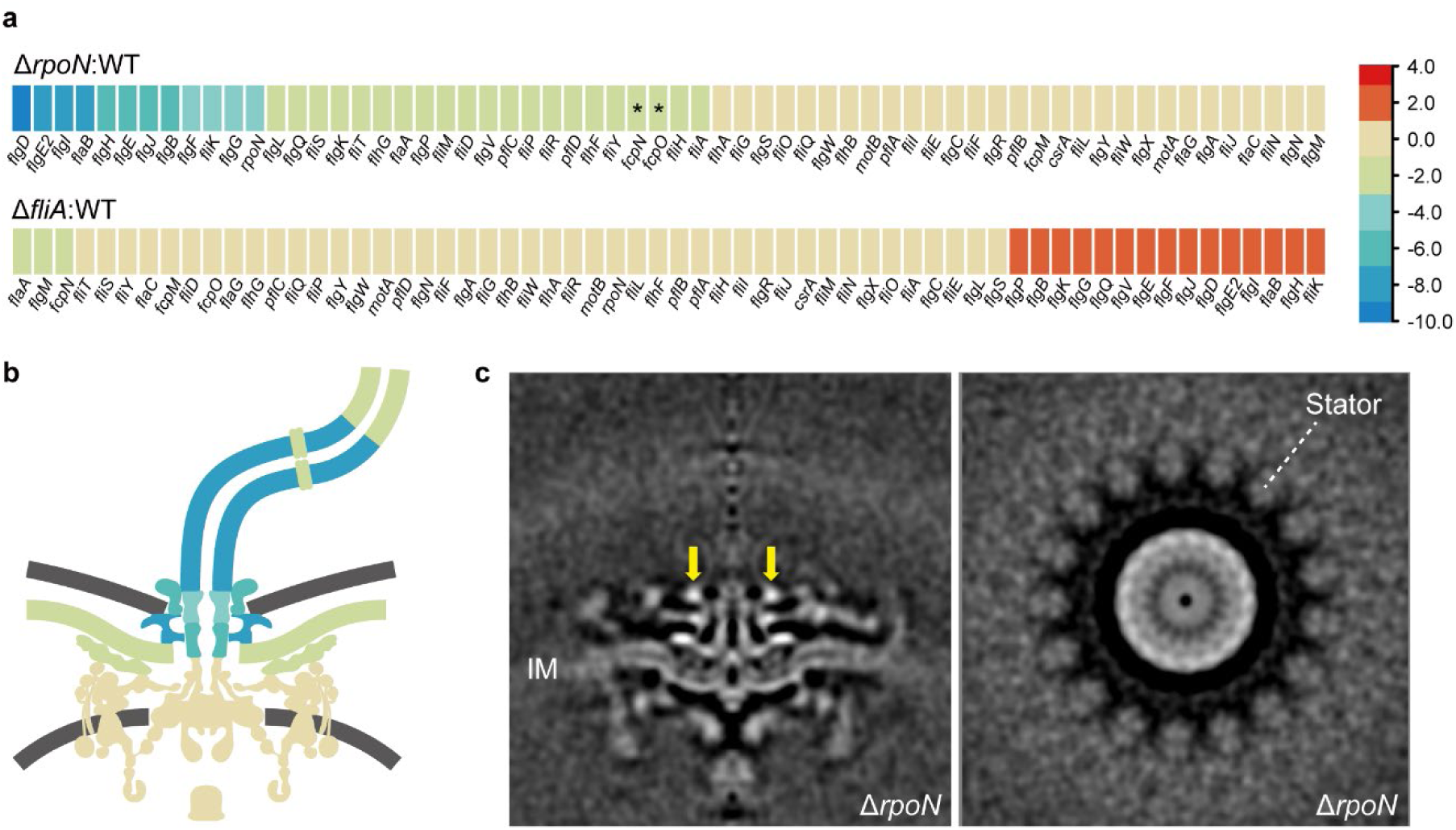
The assembly order of *C. jejuni* motor. **a**, Relative expression level of flagellar genes in two *C. jejuni* mutants Δ*rpoN* and Δ*fliA* compared to wild type, revealed by RNA-seq. The colors in the heatmaps are defined by log_2_(fold change) of FPKM values of mutant compared to wild type as indicated on the right and Methods. Asterisks (*) denote that the expression levels of *fcpN* and *fcpO* are very low in all nine samples of RNA-seq. **b**, Cartoon of motor structure from wild-type *C. jejuni* with components colored by log_2_(fold change) values of Δ*rpoN* mutant compared to wild type as indicated in **a**. **c,** Central section (left) and cross-section (right) of motor structure from Δ*rpoN* mutant revealed the early assembly of inner-membrane proximal scaffolds and stator complexes before the rod. A transient periplasmic ring above the E-ring is highlighted by yellow arrow.

As RpoN is an early checkpoint during flagellar assembly, the motor structure of the Δ*rpoN* mutant was examined by cryo-ET to gain detail for complex motor intermediates. Consistent with the RNA-seq data, the flagellar type III secretion system, C-ring, MS-ring, and inner-membrane proximal scaffolds are clearly visible in the Δ*rpoN* mutant, which lacks a rod and other periplasmic and extracellular structures (Fig. 5c). Importantly, the stator complexes seem to partially occupy the Δ*rpoN* motor, with less-resolved densities and incomplete FliL rings also present (Fig. 5c). The early assembly of these structures is also supported by a recent cryo-ET study showing flagellar intermediate structures containing the “MS-complex”, cage, and stator complexes without the rod in wild-type *C. jejuni* and *H. pylori* ^55^.

A novel density, a continuous ring without clear symmetry, also exists above the MS-ring and E-ring in the Δ*rpoN* mutant (Fig. 5c). This is likely a transient periplasmic ring that has not been reported previously in any species or in the fully assembled motor. Though its composition remains to be determined, it likely assists assembly of the rest of the basal body.

### Inner-membrane proximal scaffolds likely existed in the ancestral flagellar motor of *Campylobacterota*

To address the origin of the intricate inner membrane-anchored scaffold network in *C. jejuni*, we first analyzed the distribution of scaffolding proteins in representative species of the phylum *Campylobacterota*. This phylum is ecologically diverse, spanning from deep-sea hydrothermal vents to terrestrial environments and various hosts ^31,56^. Homologs of FlgY and PflA are present in all species of this phylum except six species without a flagellar gene set; PflB homologs are missing from species that either lack a flagellar gene set or belong to the genus *Nitratiruptor* but are present in all the remaining species; homologs of PflD and the three unfused proteins FcpM/FcpN/FcpO exist in most species of this phylum (59 out of 82 species) (Supplementary Table 4) ^31^. Importantly, the genes that flank *flgY*, *pflA*, *pflB, pflD,* and *fcpMNO* are also conserved across the genomes (Fig. 6a, Extended Data Fig. 9 and 10).

**Fig. 6.**
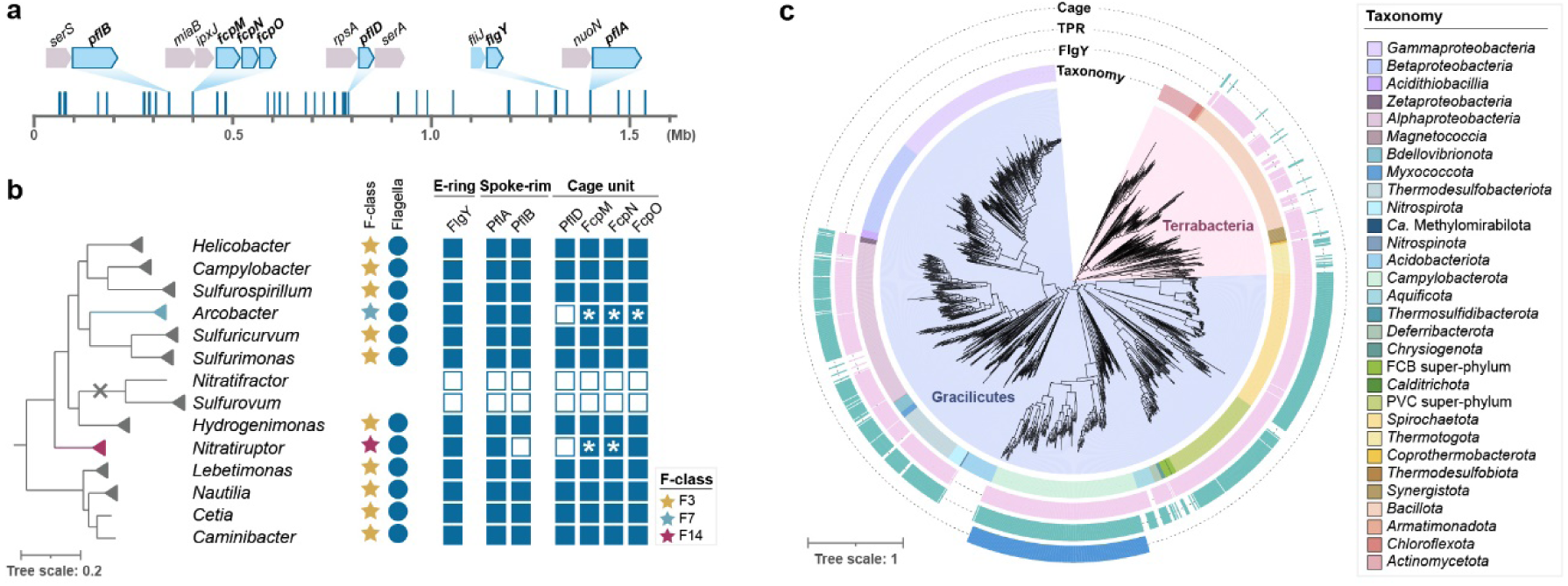
Evolution of inner-membrane proximal scaffolding proteins. **a**, The distribution of flagellar genes (represented by blue lines) in linearized genome of *C. jejuni* 81-176 and the conserved gene order of *flgY*, *pflA*, *pflB*, *pflD*, *fcpMNO* in the phylum *Campylobacterota.* Details of gene order for each species of *Campylobacterota* is provided in Extended Data Fig. 8-9. **b**, The co-occurrence of inner-membrane proximal scaffolds and the F3 chemosensory class in *Campylobacterota.* Left: the phylogenetic tree of *Campylobacterota* taken from (ref. 30), with *Wolinella* genus grouped into its close relative *Helicobacter* clade and two branches highlighted for *Arcobacter* (blue) and *Nitratiruptor* (red) that have non-F3 chemosensory class; Middle: the chemosensory F class controlling flagellar motility and the presence of flagellar gene sets across genera; Right: the presence (solid squares) and absence (hollow squares) of components of inner-membrane proximal scaffolds. Asterisks (*) denote fusion of genes *fcpMNO* or *fcpMN* in *Arcobacter* and *Nitratiruptor*, respectively. **c**, Phylogenetic distribution of proteins constituting E-ring, spokes, and cage in the *Bacteria* domain. The maximum likelihood tree of 1,365 flagellated species based on concatenated 120 marker proteins is rooted between *Terrabacteria* (peach) and *Gracilicutes* (lavender). The innermost ring is color-coded to represent the taxonomy of each species with the taxon listed on the right, while the outer rings feature colored blocks to indicate the presence of FlgY, TPR repeat proteins in proximity to flagellar gene cluster, and cage proteins.

Next, the presence of homologs was mapped to the phylogenetic tree of *Campylobacterota* ^31^. In the context of species evolution, information for the chemosensory system and the whole flagellar gene set was also added to provide a genomic background for our proteins of interest ^31^ (Fig. 6b, Extended Data Fig. 9 and 10). Clearly, species that have all seven proteins (FlgY, PflA, PflB, PflD, and unfused FcpM/FcpN/FcpO) also have the F3 class of the chemosensory system, which co-evolved with the ancestral flagellar gene set in this phylum, based on our previous studies ^31^ (Fig. 6b, Extended Data Fig. 9 and 10). To examine the association of FcpM/FcpN/FcpO homologs with flagella or T4P, genes that encode T4P components were searched for in all representative genomes of *Campylobacterota.* The T4P gene set can be identified in half of the species of this phylum, but their species distribution pattern is distinct from FcpM/FcpN/FcpO homologs (Extended Data Fig. 10 and Supplementary Table 4). In particular, all T4P gene clusters have the same gene order *pilMNOPQ*, and most have *pilM* and *pilN* fused into one gene (Extended Data Fig. 10). By contrast, *fcpMNO* is an independent operon with conserved upstream gene *miaB,* and this operon is present in all species with a flagellar gene set, regardless of the presence of the T4P gene set (Extended Data Fig. 10). Hence, based on their strict co-occurrence with the flagellar gene set, rather than T4P, in *Campylobacterota* genomes, we conclude that FcpM, FcpN, and FcpO are flagellar proteins in this phylum.

Altogether, inner-membrane proximal scaffold components are conserved in *Campylobacterota*. Structures composed of these components are likely present in other species of this phylum that inherited the ancestral flagellar gene set and F3 chemosensory class ^31^. In addition, these scaffolds likely existed in the common ancestor of *Campylobacterota*.

### The E-ring and spokes are ancient and widespread in the *Bacteria* domain

We then extended our phylogenetic analyses of scaffolding proteins to 2,638 representative species of the *Bacteria* domain, covering a great diversity of phyla/superphyla. Among the 1,365 species with a flagellar gene set, 66% have a FlgY homolog, while most species without FlgY come from two lineages: β- and γ-proteobacteria (Fig. 6c and Supplementary Table 5). Unlike FlgY with its conserved ARM-like domain and relatively conserved gene order in various genomes (Extended Data Fig. 11), neither PflA nor PflB is conserved at the sequence level. Lack of sequence conservation is a feature of TPR repeats, hindering their identification by homology search ^57^. Thus, we considered proteins with many TPR repeats that lie within or in proximity to flagellar gene clusters as potential PflA and PflB homologs. Among the species with a set of flagellar genes, approximately 52% contain TPR repeat proteins possibly related to flagella, which may be an underestimate due to the criterion of proximity to flagellar genes (Fig. 6c and Supplementary Table 5). In addition, most of these genomes have only one copy of a TPR repeat protein in proximity to flagellar genes. Structurally, this single-copy TPR repeat protein is more similar to PflA than PflB because it lacks the defining feature of PflB here, which is the presence of both a transmembrane motif and antiparallel β-sheet pair in the middle of the 3D structure (Extended Data Fig. 12).

The wide distribution of FlgY and potential PflA homologs in flagellated bacterial species across phyla suggests that the E-ring and spokes are a common structure in flagellar motors outside β- and γ-proteobacteria. Notably, several flagellar proteins in species of distantly related α-proteobacteria and *Spirochaetota* are FlgY or PflAB homologs based on structural similarity (Extended Data Fig. 11 and 12). This homologous relationship was not previously known due to low sequence homology, but the periplasmic location, protein-protein interaction, and functional relationship with stator complexes of these proteins previously reported ^23,24,58,59^ agree with our findings for FlgY and PflAB. In addition, homologs for FlgY and PflA exist in the genome of *C. crescentus* (CC_2059 and CC_2058), with the corresponding genes adjacent to each other (Extended Data Fig. 11 and 12), confirming that the “E-ring” in isolated basal bodies of this species consists of FlgY and PflA ^40,41^.

In contrast to the wide presence of FlgY and potential PflA homologs across different bacterial phyla, homologs of FcpM, FcpN, FcpO, and PflD were only found in *Campylobacterota* (Fig. 6c). Thus, the cage is a unique structure in flagellated species of *Campylobacterota*, likely evolved in the common ancestor of this phylum. Remarkably, the ancestral motor of *Campylobacterota* likely had multiple components from T4P, including the PflA_AMIN_ and PflD_PilQ-N1_ domains from PilQ, PflD_PilP-loop_ from PilP, and FcpMNO from PilMNO (Extended Data Fig. 1c, 5a, 6b-c, 13b-c). In addition, both PflA and PflB are similar to another T4P protein, PilF ^60^, as all three proteins feature multiple TPR repeats, though PflA and PflB have additional β-sheets and more TPR repeats than PilF (Extended Data Fig. 13a).

Finally, the wide presence of FlgY and potential PflA homologs in flagellated species of both Terrabacteria (including monoderm and atypical diderm lineages) and Gracilicutes (including most diderm lineages) suggests that the E-ring and spokes likely have an ancient origin (Fig. 6c). A recent study suggested that the root of the bacterial phylogenetic tree is between Terrabacteria and Gracilicutes, and the last bacterial common ancestor was a flagellated, rod-shaped diderm organism ^2^.

## Discussion

As the first discovered rotary nanomachine, the flagellar motor attracted scientists from diverse disciplines to study its assembly and mechanisms, focusing mainly on the *E. coli* model. Here, we developed a model for one of the most complex motors in unprecedented detail, revealing its architecture, assembly, and rotation (Supplementary Video 3). As accessory scaffolds are not present in the classical model, our work here also provides new mechanisms for motor assembly and function. For example, the early assembly of stator complexes along with scaffolds in both *C. jejuni* and *B. burgdorferi* ^53^ contrasts sharply with the late incorporation of stator complexes in the *E. coli* model ^14^. In addition, the lack of a peptidoglycan layer within the *C. jejuni* motor suggests an activation mechanism different from that of the *E*. *coli* model, which depends on the binding of MotB to the peptidoglycan layer ^61^. Furthermore, we showed that the majority of motor structures imaged by cryo-ET to date have a fixed number of discernable stator complexes and additional periplasmic scaffolds (Extended Data Fig. 14 and Supplementary Table 1). It is possible that many species with additional periplasmic scaffolds do not undergo dynamic exchange of stator complexes or do so with a much lower frequency of exchange than in the *E. coli* model.

The origin and evolution of the flagellum were publicly debated in 2005 due to its structural complexity, and the exact origin of the flagellum and composition of the ancestral motor remain elusive ^62,63^. Our results suggest that the E-ring and spokes, made of ancient modules such as ARM and TPR repeats ^64,65^, are widely present in modern species and likely evolved in the motor of the last bacterial common ancestor. Thus, the classical model with the simplest motor, also a product of horizontal gene transfer, cannot serve as the prototype of the ancestral motor. Interestingly, the ancestral motor of the phylum *Campylobacterota* likely recruited part of the T4P structure to generate spokes and cage units that enclose the stator complexes for flagellar motility (Supplementary Video 4). This unique evolutionary event provides a good example of exaptation: the recruitment of elements that initially evolved for other functions in other cellular structures ^66^.

We recently reported that other accessory proteins and regulators were likely present in the ancestral motor of the phylum *Campylobacterota* ^31^. By contrast, the later, simpler motor structures in two lineages of this phylum resulted from gene loss or horizontal gene transfer after loss of the whole ancestral gene set ^31^. Therefore, we propose that the evolution of complex flagellar motors is complicated requiring robust bacterial phylogeny to track and disentangle. Finally, the evolution of complex motors cannot be attributed simply to host adaptation, which is biased by studies on bacterial pathogens. For example, many modern species of *Campylobacterota* have genes that encode all known periplasmic scaffolds and live in deep-sea hydrothermal vents, on the ocean surface, or in terrestrial environments ^31^. These modern species, including the pathogens *C. jejuni* and *H. pylori*, likely inherited the ancestral flagellar gene set rather than acquiring scaffolding genes as a recent adaptation during host association.

## Materials and methods

### Bacterial strains and culture conditions

The list of strains, plasmids, and related antibiotics are summarized in Supplementary Table 6 and primers are listed in Supplementary Table 7. The *C. jejuni* 81-176 wild-type (WT) and mutant strains were routinely grown on blood agar plates (Trypticase soy agar supplemented with 5% sheep blood) at 37°C in BACTROX-2 microaerobic workstation (SHELLAB, USA) equilibrated to a 5% O_2_ and 10% CO_2_ atmosphere. For liquid cultures, *C. jejuni* strains were grown in Brain Hart Infusion (BHI) medium. The *C. jejuni* mutants were selected on Brucella broth agar plates supplemented with antibiotics as indicated below. *E. coli* was grown on LB medium or agar plates at 37°C under aerobic conditions. The selection medium contained antibiotics at the following concentrations: chloramphenicol: 50 µg ml^-1^ for *E. coli* and 10 µg ml^-1^ for *C. jejuni*; kanamycin: 50 µg ml^-1^; apramycin: 50 µg ml^-1^; ampicillin: 100 µg ml^-1^. All *C. jejuni* strains were stored at −80°C in BHI medium with 30% glycerol, and *E. coli* strains were stored at −80°C in LB medium with 15% glycerol.

### *C. jejuni* mutant construction and complementation

*C. jejuni* knockout mutant strains were constructed by the gene insertion or replacement strategy, in which an antibiotic resistance cassette was inserted into the open reading frame (ORF) of the target gene as previously described ^67^. The upstream and downstream regions of the target gene (approximately 1 kb each fragment) were PCR amplified and introduced a BamHI or EcoRI restriction enzyme cutting site in the middle. The two fragments were fused into the linearized pBluescript II SK plasmid following the Gibson assembly protocol^68^. The resulting plasmid was digested with BamHI or EcoRI enzyme, then a kanamycin or apramycin gene cassette was inserted by T4 ligase. The recombination plasmids were transformed to *E. coli* DH5α and transformants were selected on LB plates containing kanamycin or apramycin. All plasmids were verified by DNA sequencing and were naturally transformed into *C. jejuni* for gene allelic exchange. The transformants were selected on Brucella broth agar plates with kanamycin or apramycin. The mutation was confirmed by PCR analysis and DNA sequencing.

The *C. jejuni* gene knockout mutants were complemented by inserting the wild-type copy of the target gene into the *hsdR* locus with a chloramphenicol resistance cassette and a 3×FLAG tag fused to the target gene as previously described ^33^. The complemented mutants were selected on Brucella broth agar plates with chloramphenicol and kanamycin/apramycin. PCR tests were used to verify the recombinant gene regions of all constructs.

### Soft agar motility assay

The *C. jejuni* strains were incubated on blood agar plates for 24 h under microaerobic conditions at 37°C. A sterilized tip was used to dip into the colony, which was then stabbed into semisolid Brucella broth plates with 0.3% agar. The plates were incubated microaerobically (85% N_2_, 10% CO_2_, 5% O_2_) at 37°C for 20-36 h.

### Immunoprecipitation of interacting proteins and LC-MS/MS analysis

*C. jejuni* strains expressing a 3×FLAG-tagged version of the different proteins were grown on blood agar, resuspended in phosphate-buffer saline (PBS), pelleted at 6,000 rpm, and then resuspended in 2 ml of Tris-buffered saline (TBS), 1% Triton X-100, and 0.5 mM MgSO_4_ containing protease inhibitors and 10 µg ml^-1^ DNase. After lysis by sonication, cell debris were removed by centrifugation at 14,000 rpm, supernatants were recovered, and immunoprecipitation (IP) of 3×FLAG-bait protein was performed using anti-FLAG M2 affinity gel following the manufacturer recommendations. Bound proteins were eluted twice by acid elution with 40 µl of 0.1 M glycine-HCl buffer at pH 3.5. All the elution products were pooled and loaded onto a 10% SDS-PAGE gel for LC-MS/MS. The identification of IP products by LC-MS/MS was conducted as previously described ^69^.

### Cryo-ET sample preparation

*C. jejuni* strains were streaked on Tryptic Soy Agar (TSA) plate and grown at 37°C for overnight. Then, bacteria were harvested and inoculated into BHI broth to grow at 37°C for 5 h. To grow bacteria under microaerobic condition, bacterial plate and cultures were put in a jar with CampyGen^TM^ 2.5L (Thermo Fisher Scientific). 50 µg ml^-1^ of kanamycin was added into TSA plate and BHI medium to grow all mutant strains. For cryo-ET sample preparation, bacterial pellets were prepared by centrifugation with 1,500 ×g for 10 min and resuspended with PBS to a final OD_600_ of 1.0. BSA coated gold tracer solution with 10 nm particle size (Aurion) was then added to the bacterial resuspension at a ratio of 1:1 (V/V). 5 μl of the mixture was deposited on the carbon side of discharged cryo-EM grids (Quantifoil, Cu) at room temperature. Filter paper (Whatman^TM^) was used to blot on the backs of grids for almost 4 s, and cryo-EM grids were immediately plunged into liquid ethane and propane mixture using a manual gravity plunger as described previously ^27^. EM GP2 plunger (Leica) was also used to prepare cryo-EM samples. Briefly, GP2 environmental chambers were set to 25°C and 95% humidity. 5 μl of the mixture were applied to the carbon side of the discharged cryo-EM grids (Quantifoil, Cu). The grids were then blotted for 6 s and immediately plunge frozen in the liquid ethane and propane mixture.

### Cryo-ET data collection

Frozen-hydrated specimens of *C. jejuni* were imaged below −180°C using Titan Krios G2 300 kV transmission electron microscope (Thermo Fisher Scientific) equipped with a field emission gun, K3 direct detection camera (Gatan), and GIF BioQuantum imaging Filter (Gatan). The low-dose mode in SerialEM software ^70^ was used to record tilt series images at 42,000× magnification with a physical pixel size of 2.148 Å. The tilt series images from *ΔrpoN* mutant were recorded below −180°C using Glacios 200 kV transmission electron microscope (Thermo Fisher Scientific) equipped with a field emission gun with K3 direct detector (Gatan). The images were recorded at 17,500× magnification with a physical pixel size of 2.466 Å using SerialEM software ^70^. The angle of the tilt series ranged from −48° to +48° in increments of 3°, and the stage was tilted with the dose-symmetric scheme in FastTomo script ^71^. The total accumulated electron dose was ∼65e^-^/Å^2^ distributed across 33 images in the tilt series, and 10 frames of each image were recorded using the dose fraction mode in SerialEM. Parameters of cryo-ET data aquation were summarized in Supplementary Table 8.

### Cryo-ET data analysis

MotionCor2^72^ was used to correct the image drifting induced by the electron beam during the data aquation. Then, IMOD software was used to create image stacks and align all images in each tilt series by tracking with the 10 nm fiducial bead ^73,74^. For this tilt series alignment step, almost 10 fiducial beads in each image were tracked across the tilt series. For tilt series with nonsufficient numbers of fiducial beads, the patch tracking in IMOD was used for the alignment. Gctf ^75^ was used to estimate defocus for all images in the aligned tilt series, and contrast transfer function (CTF) was corrected using the ctfphaseflip function in IMOD ^76^. The binvol function in IMOD was used to generate 6× binning of the aligned stacks. 6× binned tomograms with Simultaneous Iterative Reconstruction Technique (SIRT) were reconstructed for the particle picking, and tomograms with weighted back projection (WBP) were reconstructed for the subtomogram averaging. Both SIRT and WBP reconstruction was done by Tomo3D ^77^. In total, 1,601 tomograms were analyzed (Supplementary Table 8).

### In-situ flagellar motor structure analysis by subtomogram averaging

Flagellar motors at the bacterial pole region were manually picked using the tomopick command in i3 software package ^78–80^, and the total numbers of flagellar motors for the subtomogram averaging are shown in Supplementary Table 8. I3 software package was used to align particles and determine subtomogram averaged structure in 6× binned tomograms. For further structural analysis, subtomograms were extracted based on the aligned positions of flagellar motors in the 6× binned tomograms. Then, 2× and 4× binned subtomograms were generated using the binvol function in IMOD. The 3D classification was performed to remove bad particles and determine C17 symmetrical structures in averaged structures that were used for the refinement. The C17-symmetry expansion was applied for averaged structures in 4× binned subtomograms to increase the particle numbers of flagellar motors. Then, averaged structures were refined using the 2× binned subtomograms. For the focused refined structures, local areas of the motor structures in un-binned subtomograms were extracted and refined to determine higher resolution structures. The resolution was estimated by Fourier Shell Correlation (FSC).

### Molecular modeling of the *C. jejuni* flagellar motor

The atomic models of FlgY homodimer, PflA and PflB complex, FcpN/FcpO heterodimer, and FliL ring were predicted by AlphaFold3 ^81^. For the PflA and PflB complex, the predicted structure was used as template, and then the N-terminal domain of PflA and PflB were replaced with in vitro structure that was determined by single particle analysis of cryo-EM. Predicted MS-ring, FlgY, PflC, PflD, and FcpMNO complexes were used for the pseudo-atomic model. For the stator units, the published structure of MotA complex (PDB: 6ykm) ^3^ and predicted MotB model in *B. burgdorferi* flagellar motor ^82^ were used for the modeling. Cryo-EM structures of LP-ring (PDB: 7cbl) ^4^ and C-ring (PDB: 8uox) ^7^ were also used for the modeling. Based on the pseudo-atomic model of *C. jejuni* flagellar motor, UCSF ChimeraX and Blender were used to create movies based on the cryo-ET analysis of in situ structures of the motors.

### Protein expression and purification

For recombinant protein expression in *E. coli* BL21 (DE3), FlgY and its truncated variants, as well as PflA_16-788_ were cloned into pET22b vector which contains an N-terminal *pelB* signal peptide for periplasmic targeting and a C-terminal 6×His tag. PflA_16-788_ and PflB_113- 820_ were cloned into pET26b, also featuring an N-terminal *pelB* signal peptide for periplasmic localization but with a C-terminal Strep-tag instead. All constructs were confirmed by DNA sequencing.

*E. coli* BL21 (DE3) carrying plasmids encoding FlgY and its truncation variants and PflA_16-788_-Strep construct were grown in 1 L LB medium and induced with 0.2 mM isopropyl-β-d-thiogalactopyranoside (IPTG) at OD_600_ of ∼0.8. After growing for 16 h at 20°C, the cell was harvested and resuspended in high salt lysis buffer (20 mM Tris-HCl buffer at pH 8.0 and 500 mM NaCl) and lysed by high-pressure cell crusher (Union-Biotech). The lysates were centrifuged at 17,000 ×g for 50 min and the supernatants were loaded onto Ni-NTA resin (Qiagen) or Strep-affinity beads (IBA Lifesciences). After washing with high salt lysis buffer, the proteins were eluted with 250 mM imidazole and further purified by gel filtration chromatography (Superdex 200 Increase 10/300 GL, GE Healthcare) with lysis buffer (20 mM Tris-HCl buffer at pH 8.0 and 300 mM NaCl).

For PflA_16-788_ and PflB_113-820_ complex protein purification, the growth medium was changed from LB to TB and the lysis buffer composition was adjusted to include 300 mM NaCl. All other purification steps remained unchanged. For MST experiments, all buffers were replaced with HEPES buffer at pH 7.5.

### Cryo-EM sample preparation and data collection

Aliquots of 4 µl of the PflA_16-788_ and PflB_113-820_ complex, at a concentration of 0.4 mg ml^-1^, were applied onto glow-discharged holey Quantifoil carbon-coated grids (Cu R1.2/1.3, 300 mesh, Beijing Zhongjingkeyi Technology, Beijing, China). Following a 5 s incubation on the grids under 100% humidity, the grids were blotted for 3 s at 8 ℃ using a blot force of 1, and then plunged into liquid ethane using a Vitrobot Mark IV (Thermo Fisher Scientific). EPU software (Thermo Fisher Scientific) was employed for automated data collection on a Glacios 2 transmission electron microscope equipped with an energy filter. The data were collected using a Falcon4i camera in super-resolution mode, with a defocus range of −0.5 to −1.5 μm and at a nominal magnification of 130,000× (resulting a calibrated physical pixel size of 0.89 Å). The accumulated dose was set to 40 electrons per Å^2^. Data acquisition parameters, including exposure time, beam intensity, and drift correction settings, are detailed in Supplementary Table 9.

### Cryo-EM image processing

A total of 5,055 micrographs were acquired and imported into cryoSPARC ^83^. Drift correction and dose-weighting were conducted using MotionCor2 ^72^. CTF parameters were estimated using CTF Estimation (CTFFIND4) in cryoSPARC ^83^. Subsequently, 1,265,294 particles were auto picked and extracted from 4,661 micrographs with a box size of 320 pixels for 3 rounds of 2D classification, after this cleaning step, only 161,182 particles remained for further analysis. All extracted particles were utilized for 3D classification. These particles were classified into four classes after ab-initio reconstruction and heterogeneous refinement and NU refinement, resulting in a structure at 3.55 Å resolution.

To further improve the resolution, we referred to a “seed” strategy ^84^, the raw particles were divided into 6 subgroups and combined with the “seed” particles, which was the NU refinement result. The combined particle subsets were subjected to multiple rounds of guided multi-reference 3D classification. After the optimized procedure for data processing, the dataset of PflA_16-163_ and PflB_113-820_ gave rise to the final structure at 3.23 Å resolution. A flowchart showing the data processing is shown in Extended Data Fig. 4.

### Model building, refinement, and visualization

The initial model of PflA_16-163_ and PflB_178-820_ complex was built by Cryo-Net sever ^85^, and manually adjusted in Coot ^86^. Model refinement was performed using *phenix.real_space_refine* tool within Phenix ^87^. Cryo-EM structure and model were visualized by using PyMOL and ChimeraX. The refinement statistics are summarized in Supplementary Table 9.

### Microscale thermophoresis (MST) binding assay

MST measurements were conducted on a Monolith NT.115 (NanoTemper) with 20% LED power and 40% MST power. Purified proteins (FlgY_15-172_/ FlgY_90-172_) were diluted at a concentration of 200 nM in the MST buffer containing PBS and 0.05% (w/v) Tween-20. PflA_16-788_-Strep protein was titrated from 0 to 60 μM, and then the samples were loaded into MST NT.115 standard glass capillaries for measurements. Data represent mean ± SEM of three independent measurements.

### Protein cross-linking

Purified FlgY_15-172_/ FlgY_90-172_ protein was diluted to a final concentration of 25 μM in HEPES buffer (20mM HEPES buffer at pH 7.5, 300 mM NaCl). The mixture was incubated with EGS to a final concentration of 1 mM, dissolved in anhydrous DMSO, for 30, 60 or 90 min at room temperature. The DMSO concentration was consistently maintained at 1% (V/V) of the total reaction volume. Reactions were terminated by adding 2 μl 1 M Tris-HCl buffer at pH 8.0, followed by mixing for 5 min at room temperature. Samples were subsequently boiled for 5 min and subjected to 15% SDS-PAGE analysis.

### Bacterial two hybrid (BTH) analysis

The assays were performed as described to investigate protein-protein interactions ^88^. Proteins of interest were fused to T25 or T18 fragments in plasmids pKNT25/pKT25 or pCH363/pUT18C (Supplementary Table 6). The pair of plasmids expressing fusion proteins with T18 or T25 fragments was co-transformed into *E. coli* strain BTH101 and plated on LB agar containing ampicillin (100 µg ml^-1^) and kanamycin (50 µg ml^-1^). Multiple transformants were inoculated into individual tubes with 0.5 ml of LB broth containing the same antibiotics and 0.5 mM IPTG. The cultures were incubated for 8 h at 30°C with shaking. Two microliters of each culture were spotted on LB plates containing the antibiotics, 40 µg ml^-1^ X-Gal, and 0.5 mM IPTG. The plates were incubated at 30 °C for 24 h.

### RNA extraction, RNA-seq, and quantitative real-time PCR

*C. jejuni* wild-type and Δ*rpoN*, Δ*fliA* mutants were grown separately under microaerobic conditions and harvested during the mid-exponential phase. RNA was extracted using a TIANamp RNAprep Pure Cell/Bacteria Kit. The purity and concentration of the RNA were determined by gel electrophoresis and a NanoDrop spectrophotometer (Thermo Fisher Scientific). Samples were submitted to GENEWIZ (Suzhou, China) for library construction and sequencing on an Illumina NovaSeq system. The sequencing reads were mapped to the *C. jejuni* 81-176 genome using Bowtie2 (v2.2.6). HTSeq (V0.6.1) was used to count the read numbers mapped to each gene, and then the FPKM (Fragments Per Kilo bases per Million reads) of each gene was calculated. Differential expression analysis of mutant vs. wild type was performed using the DESeq2 (V1.26.0).

For qRT-PCR analyses, total RNA was extracted as described above, followed by DNase I treatment. Then RNA was transcribed using the cDNA Master Kit (TOYOBO, Japan). Transcript levels were determined with SYBR Green Realtime PCR Master Mix (TOYOBO, Japan) in a CFX96 Connect Real-Time PCR Detection System (Bio-Rad). The cycling parameters were 95 °C for 30 s, followed by 40 cycles of 94 °C for 15 s and 60 °C for 30 s. The abundance of the *rpsQ* gene was used as an internal standard, and the relative expression levels of genes of interest were calculated using the Quantitation-Comparative CT (2−ΔΔCT) method ^89^.

### Bioinformatics analysis of flagellar inner-membrane proximal scaffolding proteins and T4P components in the phylum *Campylobacterota*

The genomes of 82 representative species of the phylum *Campylobacterota* from our previous study ^31^ were analyzed for homologs of seven flagellar scaffolding proteins (FlgY, PflA, PflB, PflD, FcpM, FcpN, and FcpO). The sequences of these proteins from *C. jejuni* 81-176 were used as queries for local BLAST search, and the best hit with an E-value < 1e^-3^ as potential homologs. Then, structural similarity of candidate homologs was confirmed using HHpred ^90^ and AlphaFold3 ^81^. All identified protein homologs are summarized in Supplementary Table 4.

To identify T4P components, MacSyFinder ^91^ was first used to identify a complete T4P gene cluster in *Campylobacter sputorum* genome (NCBI accession number: GCF_002220775.1). Then, nine core T4P components (PilM, PilN, PilO, PilP, PilQ, PilB, PilC, PilT, and PilU) from *C. sputorum* were used as queries for local BLAST searches against 82 *Campylobacterota* genomes to identify potential candidate homologs (E-value < 1e^-5^). The candidate proteins were further analyzed for domain organization using the SMART database ^92^ and for structural similarity using HHpred ^90^ and AlphaFold3 ^81^. Since T4P belongs to type IV filament (TFF) superfamily and its components share homology with other TFF systems ^93^, manual curation was performed to verify that the identified clusters belong to T4P rather than the other TFF systems. For manual curation, the identified clusters and their neighborhood genes (five genes upstream and downstream) were examined for the presence of characteristic proteins from other TFF systems based on reference ^93^. All identified T4P components are summarized in Supplementary Table 4.

To study the evolution of flagellar scaffolding proteins and T4P in *Campylobacterota*, the above identified homologs were mapped onto a species tree that also includes flagellation status and chemosensory classes of each species based on data from our recent studies ^31^. Briefly, the species tree was constructed using the UBCG pipeline ^94^, based on the sequence alignment of 92 single-copy concatenated marker proteins derived from the complete genomes of 82 *Campylobacterota* species ^31^. Six closely related *Desulfurellales* species were included as an outgroup to root the tree ^31^.

### Bioinformatics analysis of flagellar inner-membrane proximal scaffolding proteins in the *Bacteria* domain

Genomes of representative species, totaling 16,533, were downloaded from NCBI on April 3, 2023 (https://www.ncbi.nlm.nih.gov/datasets/genome/?taxon=2&reference_only=true). To address the considerable variation in the number of representative species across different taxa and ensure balanced species counts, the following criteria were applied: (1). for large groups or subgroups with more than 200 genomes, only complete genomes were selected and if the number of complete genomes exceeded 200, only one genome per genus was retained; (2). for smaller groups or subgroups with fewer than 200 genomes, all genomes were kept to ensure species diversity. Representative species were prioritized based on experimental characterization, with emphasis on model organisms listed in the COGs database ^95^, especially those whose flagellar motors have been investigated via Cryo-ET (Supplementary Table 1). Subsequently, CheckM ^96^ was used to filter genomes with completeness below 70% or contamination above 10%, resulting in a final dataset of 2,638 genomes. The information of selected 2,638 genomes was listed in Supplementary Table 5.

To determine whether the representative species has flagellar gene set, 24 core flagellar proteins (FlhA, FlhB, FliP, FliQ, FliR, FliI, FliH, FliF, MotA, MotB, FliG, FliM, FliN, FliE, FlgB, FlgC, FlgF, FlgG, FlgE, FlgD, FlgK, FlgL, FliC, and FliD) from *S. enterica* or *Bacillus subtillis* were used as queries for both BLASTP and PSI-BLAST searches ^97^. Given the presence of incomplete genomes and pseudogenes, species with at least 20 core genes were classified as possessing flagellar gene set. Consequently, 1,365 species were identified as “flagellated species” (species with flagellar gene set) and used for further phylogenetic analysis and homolog searches. A maximum likelihood (ML) phylogenetic tree was constructed for these 1,365 species using 120 markers from GTDB-Tk v2 ^98^, with the concatenated multiple sequence alignment applied to infer the tree with IQ-TREE ^99^. The best fit model, LG+R10, was selected by ModelFinder, and bootstrap was estimated based on 1,000 replications. Phylogenetic tree was visualized, annotated, and modified using iTOL v6 ^100^.

To identify FlgY homologs, comprehensive HMM homology searches ^101^ were performed following this strategy: (1). The MgtE_N Hidden Markov Model (HMM) profile (Pfam PF03448) was used to search against 1,365 flagellated species; (2). the hits obtained from the HMM search were further validated by structural analyses including SMART database ^92^, HHpred ^90^ and AlphaFold3 ^81^, but only if they were located within five genes adjacent to the flagellar gene clusters; (3) to find homologs overlooked by the HMM search, genes neighboring *fliJ* or *fliK* were also examined using SMART database, HHpred or AlphaFold3.

PflA and PflB homologs are not conserved either in sequence level or in gene order across different taxa. For some taxa with experimentally investigated homologs, these homologs were used as queries for BLAST search against the specific taxon. For example, PflAB from *C. jejuni* ^33^ were used as queries for both BLASTP and PSI-BLAST searches against *Campylobacterota*; FlcAB from *B. burgdorferi* were used for searches against *Spirochaetota* ^18,25^; and MotC from *S. meliloti* ^58^, MotK from *C. sphaeroides* ^59^, and CC_2058 from *C. crescentus* were used for searches against α-proteobacteria. For other taxa, extensive HMM homology searches were carried out according to the following criteria: (1). The TPR HMM profile (Pfam PF00515) was applied to search against the genomes of flagellated species with FlgY homologs; (2). the hits retrieved from the HMM search were further confirmed using TPRpred ^102^ and SMART database, but only if they were located within five genes adjacent to the flagellar gene clusters; (3). as TPR repeats typically consist of more than 3 consecutive motifs ^103^, candidates with less than 3 sequential motifs were discarded; (4). candidates containing functional known domains that are enriched in TPR repeats, such as glycosyltransferase, methyltransferases, acetyltransferase, sulfotransferase, metalloprotease, were further excluded.

PflD exhibits considerable sequence variability in its N-terminal loop region and its C-terminal domain that shares structural similarity with PilQ-N1 domain has no identified HMM profile in the Pfam database. Homolog searches of PflD in the phylum *Campylobacterota* were performed as described above. However, using an HMM profile built from homologs in *Campylobacterota* could introduce bias for homolog search in other phyla, thus no further bioinformatics analyses of PflD in the *Bacteria* domain were performed.

To identify homologs of cage proteins (FcpM/FcpN/FcpO), the FcpN homologous protein PilN was selected as a feature protein to locate the *fcpMNO* gene cluster. Sequence alignment was performed on all PilN proteins (COG3166) in the COG database to construct an HMM profile. To identify potential candidate homologs in 1,365 flagellated species, an HMM search was conducted (E-value < 1e^-3^). The candidate proteins were further examined for domain organization using the SMART database ^92^ and for structural similarity using HHpred ^90^ and AlphaFold3 ^81^. Additionally, since the genes of cage proteins are typically clustered together in the genome and not adjacent to any TFF family genes, gene neighborhood analysis was performed manually to confirm that the identified gene clusters belonged to *fcpMNO*.

The structures of FlgY of PflAB homologs in Extended Data Fig. 11 and 12 were predicted by AlphaFold3 ^81^. The signal peptides and transmembrane regions were predicted by SignalP 6.0 ^104^ and TMHMM 2.0 ^105^, respectively. All identified proteins are summarized in Supplementary Table 5.

## Acknowledgments

The authors would like to thank Drs. Brian Crane and Michael Lynch from Cornell University for discussion of the ARM-like domain; the High-Performance Computing Division at the South China Sea Institute of Oceanology for data analysis; Jennifer Aronson (Yale University) for editing and valuable comments on the manuscript; Yale Center for Research Computing (YCRC) for guidance and use of the research computing infrastructure; Xiaoju Li, Jing Zhu, and Zhifeng Li from Shandong University Core Facilities for Life and Environmental Sciences for their help with the Cryo-EM and MST experiments.

## Author Contributions

B.G. and J.L. conceived and designed the study. Genetic, protein interaction, RNA-seq and partial biochemical experiments were performed by X.F. with assistance from Y.L., Y.C. and M.A.B. Cryo-ET experiments and related structural analyses were performed by S.T. with assistance from H.Z. and C.H., and modeling was performed by J.M.B and S.T. Single particle cryo-EM experiments of PflAB and biochemical analyses of FlgY were performed by J.H. under the supervision of X.G. Phylogenetic analysis and homolog search were performed by S.Z. with assistance from Y.L. Partial mutants were provided by C.H. and M.L.T., and LC-MS/MS experiments were performed by M.L.T. The manuscript was written by B.G., J.L., S.T. and X.F. All authors contributed to editing the manuscript, and support the conclusions.

## Data Availability Statement

All relevant data are within the manuscript and its Supporting Information files. The cryo-ET maps of the motor in wild-type *C. jejuni*, and Δ*motA*, Δ*flgY*, Δ*pflA*, Δ*pflB*, Δ*pflC*, Δ*pflD*, and Δ*fcpMNO* mutant cells have been deposited in the Electron Microscopy Data Bank under accession codes EMD-45507, EMD-45508, EMD-49254, EMD-49256, EMD-49255, EMD-49253, EMD-49252, respectively. The cryo-EM map and atomic coordinates have been deposited in the Protein Data Bank and the Electron Microscopy Data Bank under the accession numbers 9LEQ and EMD-63032, respectively. Further details are provided in Supplementary Table 8 and 9. Raw reads of RNA-seq studies have been deposited in the Sequence Read Archive (https://www.ncbi.nlm.nih.gov/sra) under BioProject numbers PRJNA1223640. The transcriptome datasets generated in this study are summarized in Supplementary Table 3. The *Campylobacter jejuni* 81-176 reference genome (NCBI accession number: GCA_000015525.1) is available for download from NCBI (https://ftp.ncbi.nlm.nih.gov/genomes/all/GCF/000/015/525/GCF_000015525.1_ASM1552v1/).

---

**Extended Data Figure 1.**
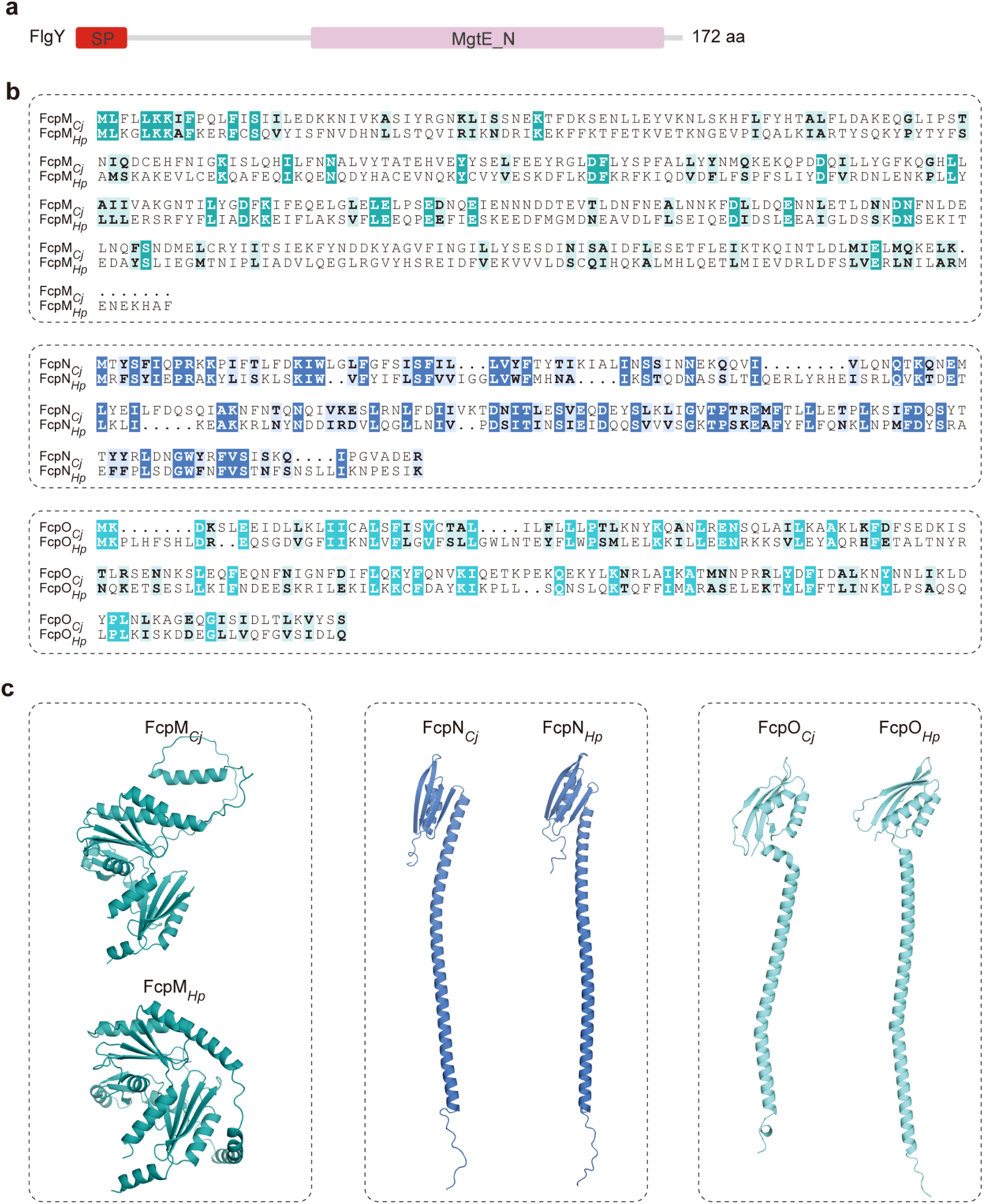
Sequence and structural analyses of FlgY and FcpMNO. **a**, SMART motif analysis of FlgY (WP_002855458.1) suggested a single peptide (SP) at its N-terminus and a MgtE_N domain at C-terminus. **b**, Sequence alignment of FcpM (WP_002868795.1), FcpN (WP_002868796.1), and FcpO (WP_002859017.1) to their homologs in *H. pylori* (WP_000911905.1, WP_001212813.1, WP_001212813.1). **c**, Comparison of AlphaFold3-predicted structures of FcpM, FcpN, and FcpO from *C. jejuni* and *H. pylori*. Sequences for structural prediction are the same as in **b**.

**Extended Data Figure 2.**
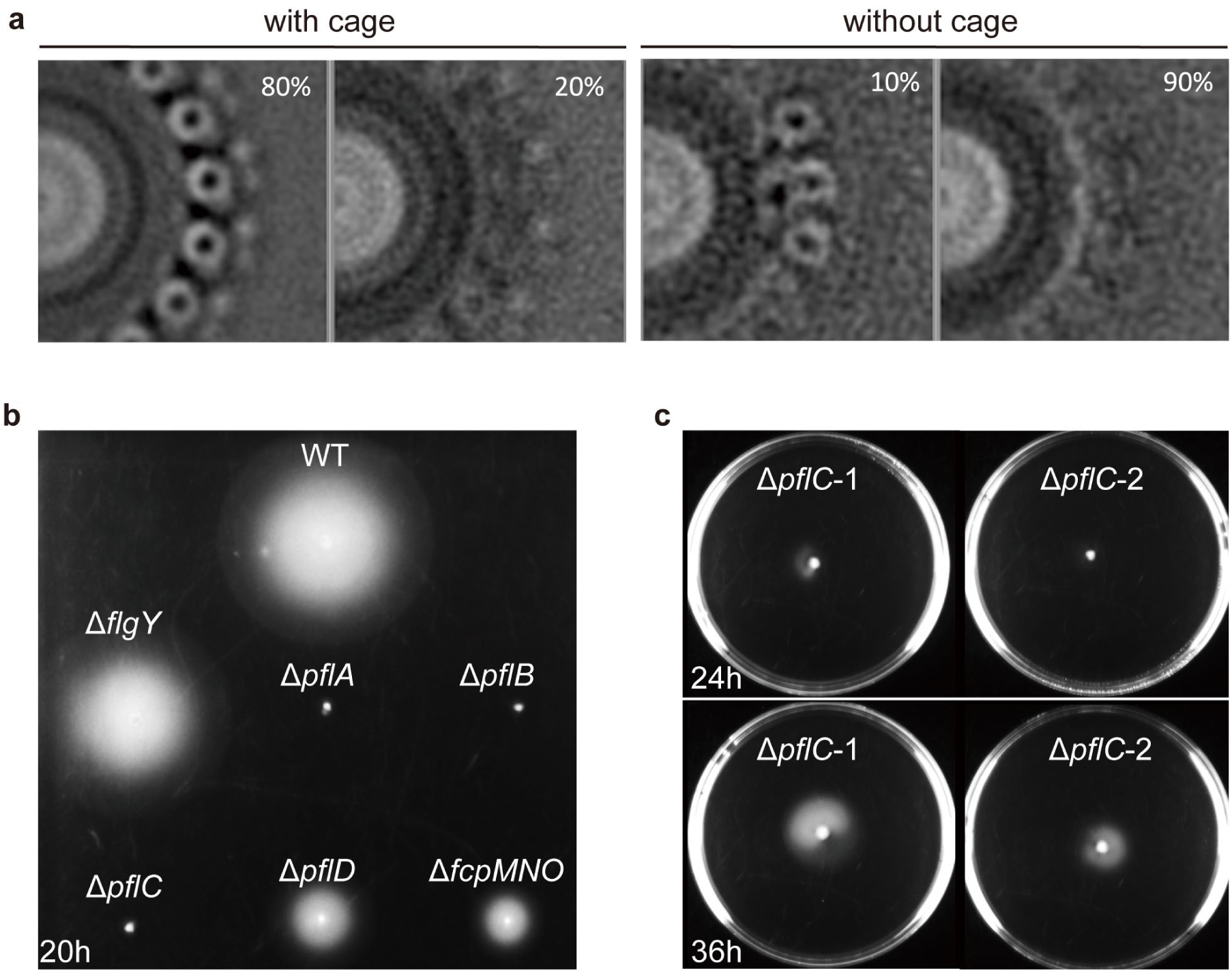
Stator occupancy and motility of WT *C. jejuni* and mutants. **a**, Cross-section of motor structure in stator ring region from WT and Δ*fcpMNO* mutant to compare stator occupancy with cage (WT) and without cage (Δ*fcpMNO* mutant). **b**, Soft agar motility assay of WT and six mutants imaged at 20 h. **c**, Soft agar motility assay of Δ*pflC* mutant imaged at 24 h and 36 h, showing reduced but not abolished motility.

**Extended Data Figure 3.**
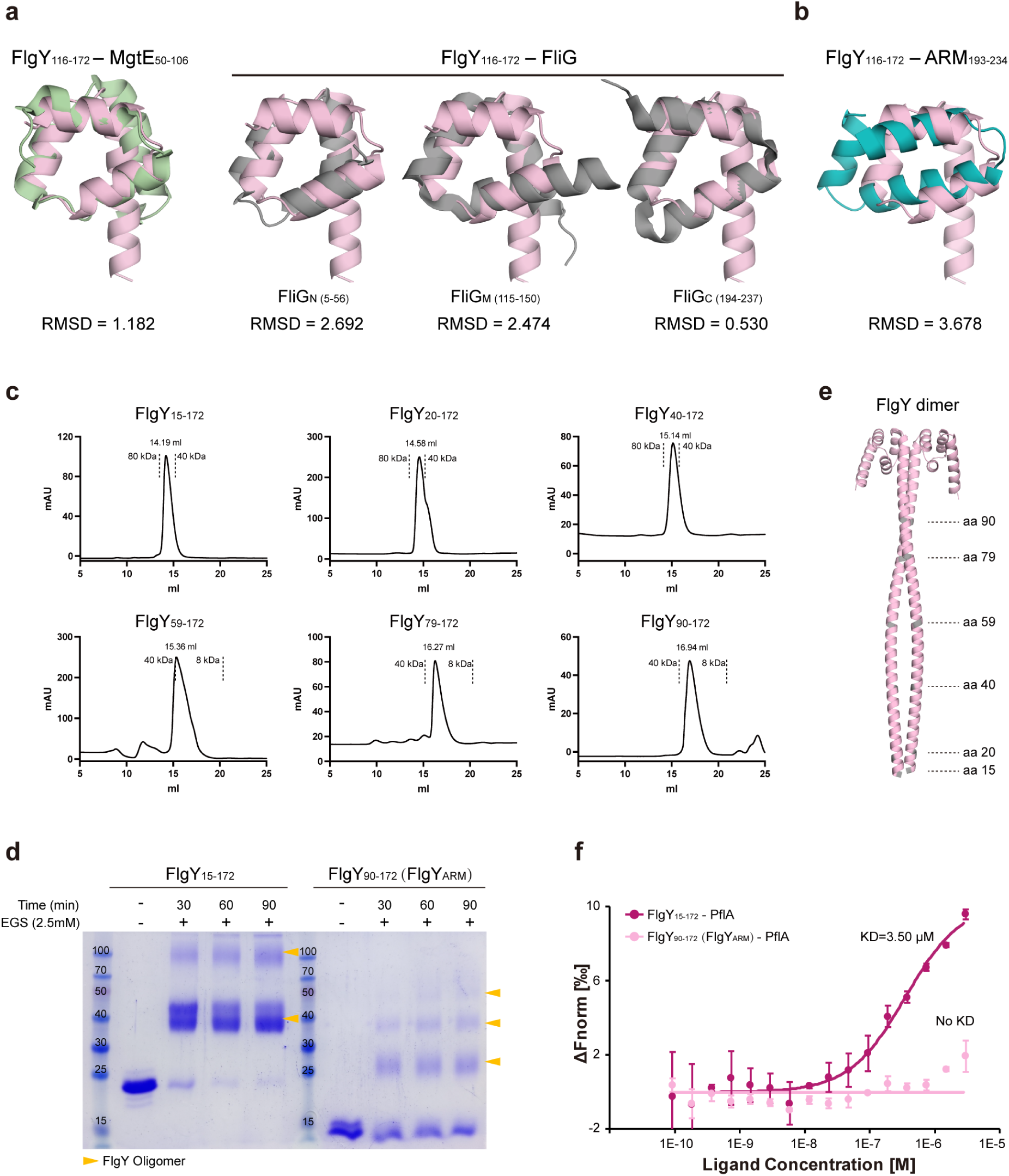
Structural analyses of FlgY. **a**, Structural overlap of FlgY C-terminal globular domain (predicted by AlphaFold3) with MgtE_N domain of MgtE (PDB: 2YVX) or three ARM-like domains of FliG (PDB: 3HJL). **b**, Structural overlap of FlgY C-terminal globular domain with canonical ARM repeat of β-catenin (PDB: 3BCT). **c**, Size-exclusion chromatography (SEC) profile of purified FlgY_15-172_ and different truncation variants. **d**, Crosslinking of purified FlgY_15-172_ and FlgY_90-172_ by adding Ethylene glycol bis(succinimidyl succinate) (EGS). Since FlgY_90-172_ is mainly composed of the ARM-like domain and forms a dimer that fits the cryo-ET map of the E-ring, we use FlgY_90-172_ to represent the dimeric FlgY_ARM_. **e**, AlphaFold3-predicted structure of FlgY dimer. **f**, MST assays of FlgY or FlgY_ARM_ with PflA to detect their interaction.

**Extended Data Figure 4.**
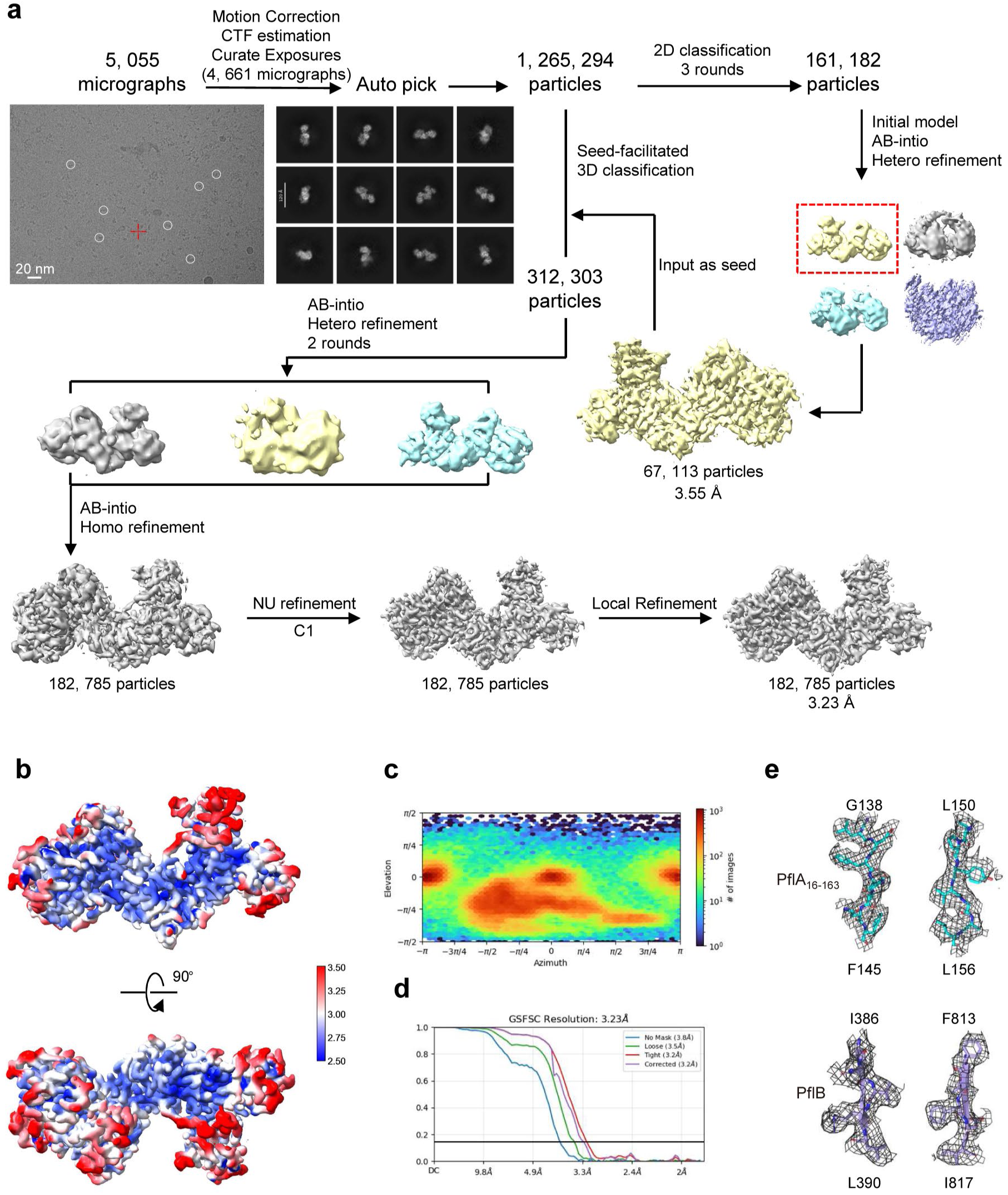
Cryo-EM structural determination of PflA_16-163_-PflB complex. **a.** Flowchart for cryo-EM data processing of the PflA_16-163_-PflB_178-820_ complex. Details can be found in Methods. The complex with C1 symmetry was reconstructed at 3.23 Å resolution. **b.** Local resolution estimations for the EM density map of the PflA_16-163_-PflB_178-820_ complex. **c.** Angular distribution of the particles used for reconstruction of the PflA_16-163_-PflB_178-820_ complex. **d.** Gold standard Fourier Shell Correlation (FSC) curves of the PflA_16-163_-PflB_178- 820_ complex. **e.** Representative Cryo-EM densities in the PflA_16-163_-PflB_178-820_ complex. The maps, shown as grey mesh, are contoured at 5–6σ.

**Extended Data Figure 5.**
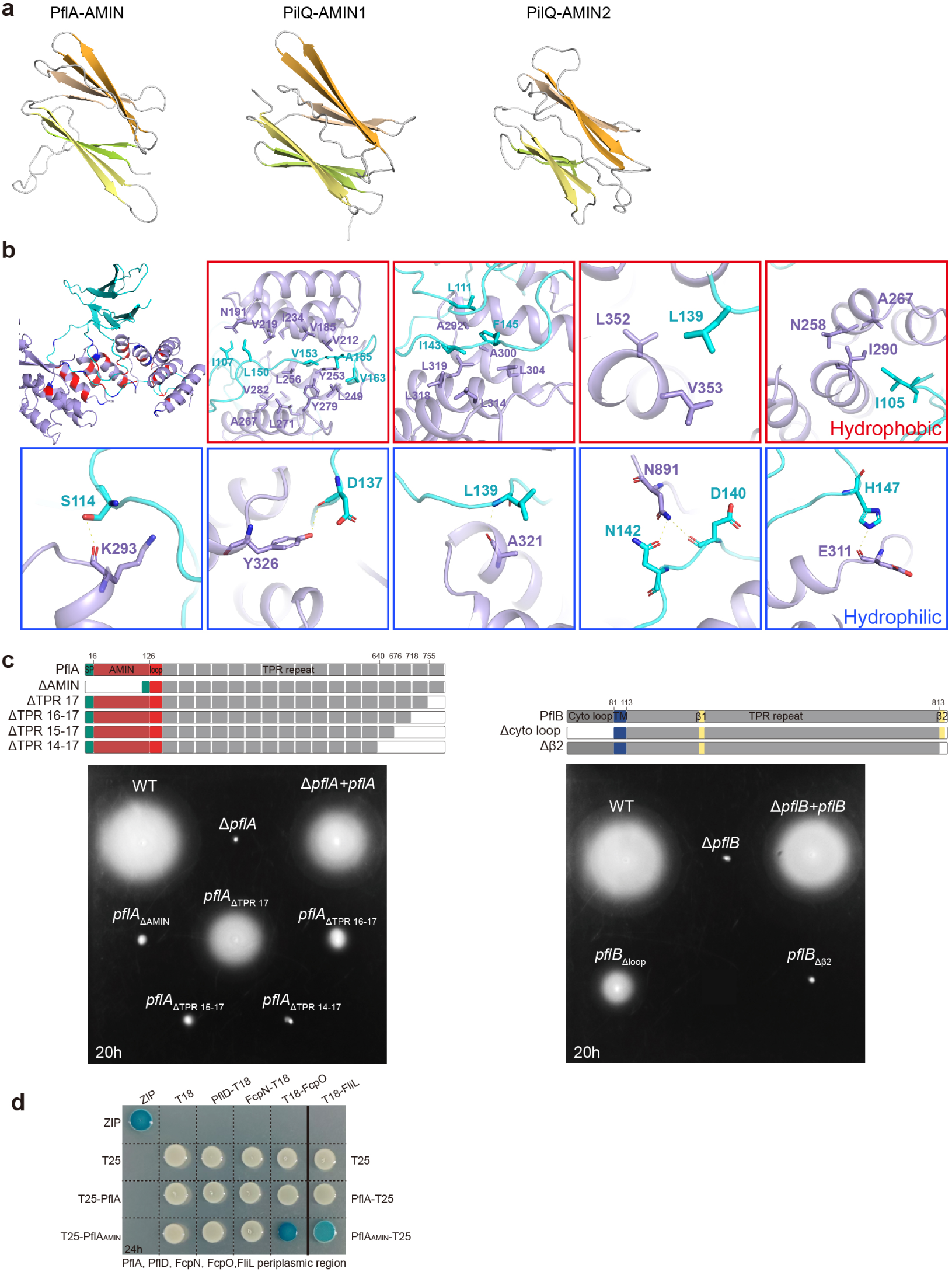
Structural analyses of PflA and PflB. **a**, Structural comparison of the β-sandwich domain of PflA with AMIN1 and AMIN2 domains of PilQ (PDB:3JC8) of T4P. **b**, Structural details of PflAB interaction interface for both hydrophobic and hydrophilic interactions. **c**, Soft agar motility assay of complementation of various truncations of PflA or PflB into Δ*pflA* or Δ*pflB* mutant, respectively. The domain organization of full-length PflA or PflB and their various truncation constructs made for soft agar motility analyses were indicated on the top. **d**, BTH analysis for the interaction of PflA or PflA_AMIN_ domain with PflD, FcpN, FcpO, and FliL. All these proteins were cloned with their periplasmic region only.

**Extended Data Figure 6.**
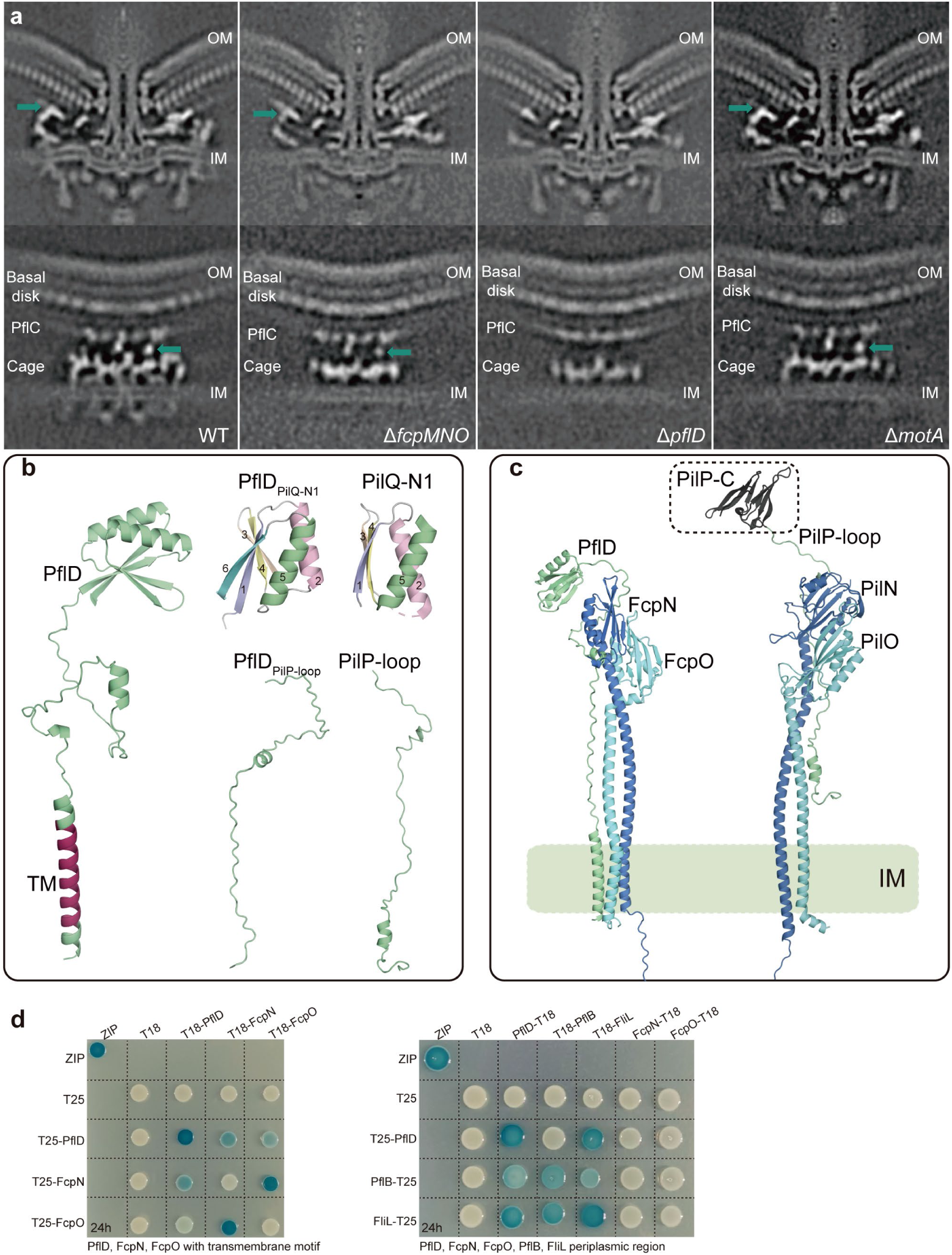
The location of PflD in motor and its interaction with FcpNO. **a**, The central cross section of the complete motor structure (top) and side view of the cage region (bottom) from WT and three mutants. The position of PflD is indicated with a green arrow. **b**, Left: AlphaFold3-predicted structure of PflD with transmembrane motif highlighted in red. Right: The C-terminal domain and middle loop region of PflD show structural similarity to the N1 domain of PilQ and the loop region of PilP, respectively. **c**, Comparison of complex structure of PflD-FcpNO predicted by AlphaFold-Multimer and PilNOP of T4P (PDB:3JC8). **d**, BTH analysis for the interaction of PflD and FcpNO or other proteins. Left: the interaction of PflD and FcpNO with their transmembrane motif and the T25 or T18 tags were cloned at the N-terminal end next to transmembrane motif; Right: the interaction of PflD and other proteins with periplasmic region only.

**Extended Data Figure 7.**
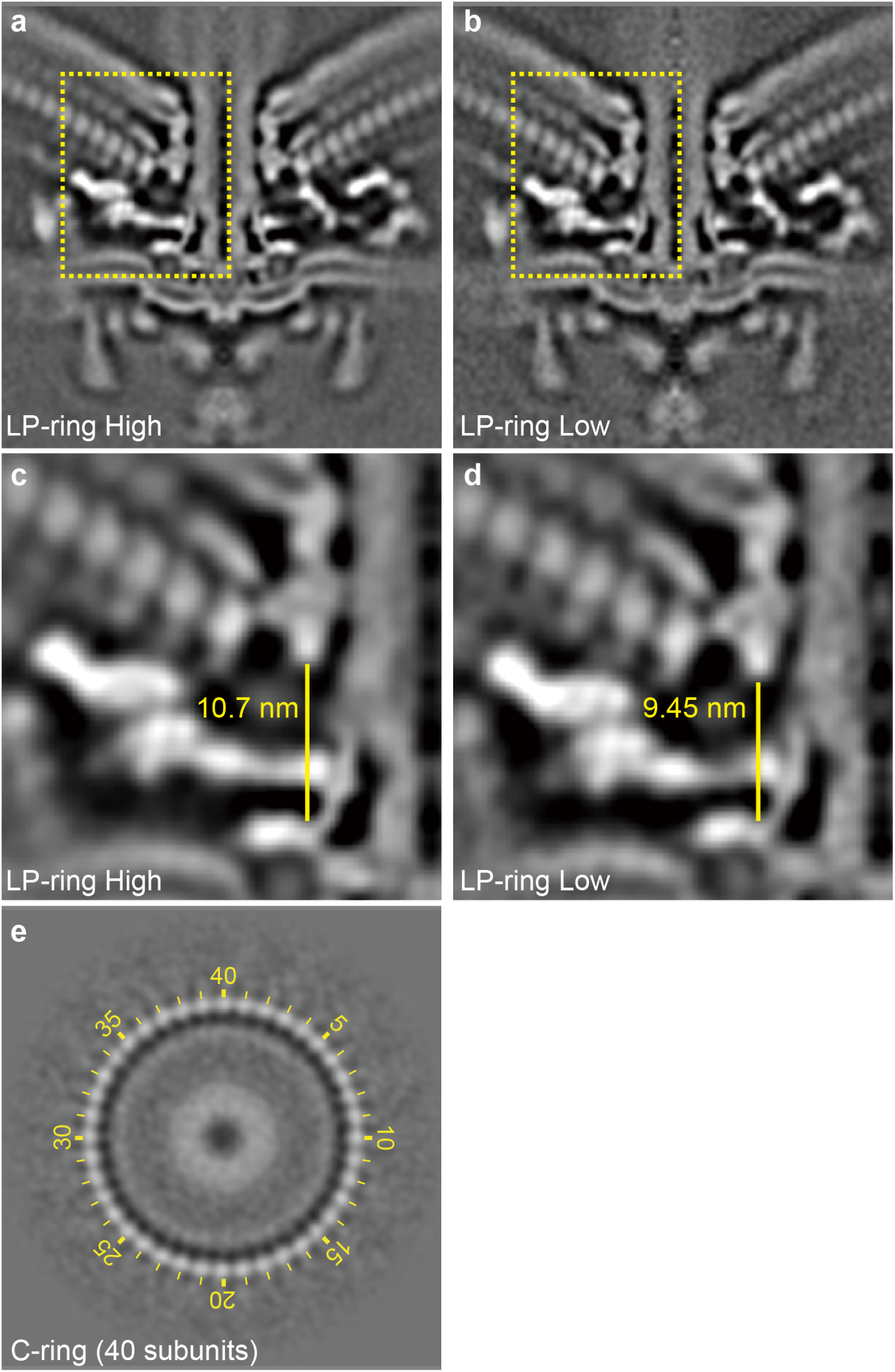
Dynamic changes and C-ring symmetry of *C. jejuni* motor. **a-b**, Central section of motor structures from WT *C. jejuni* with two distinctive heights from LP-ring to MS-ring. **c-d**, Close-up views of boxed regions in **a** and **b** with measured height from LP-ring to MS-ring, respectively. **e**, Focused classification on the C-ring reveals its 40 subunits.

**Extended Data Figure 8.**
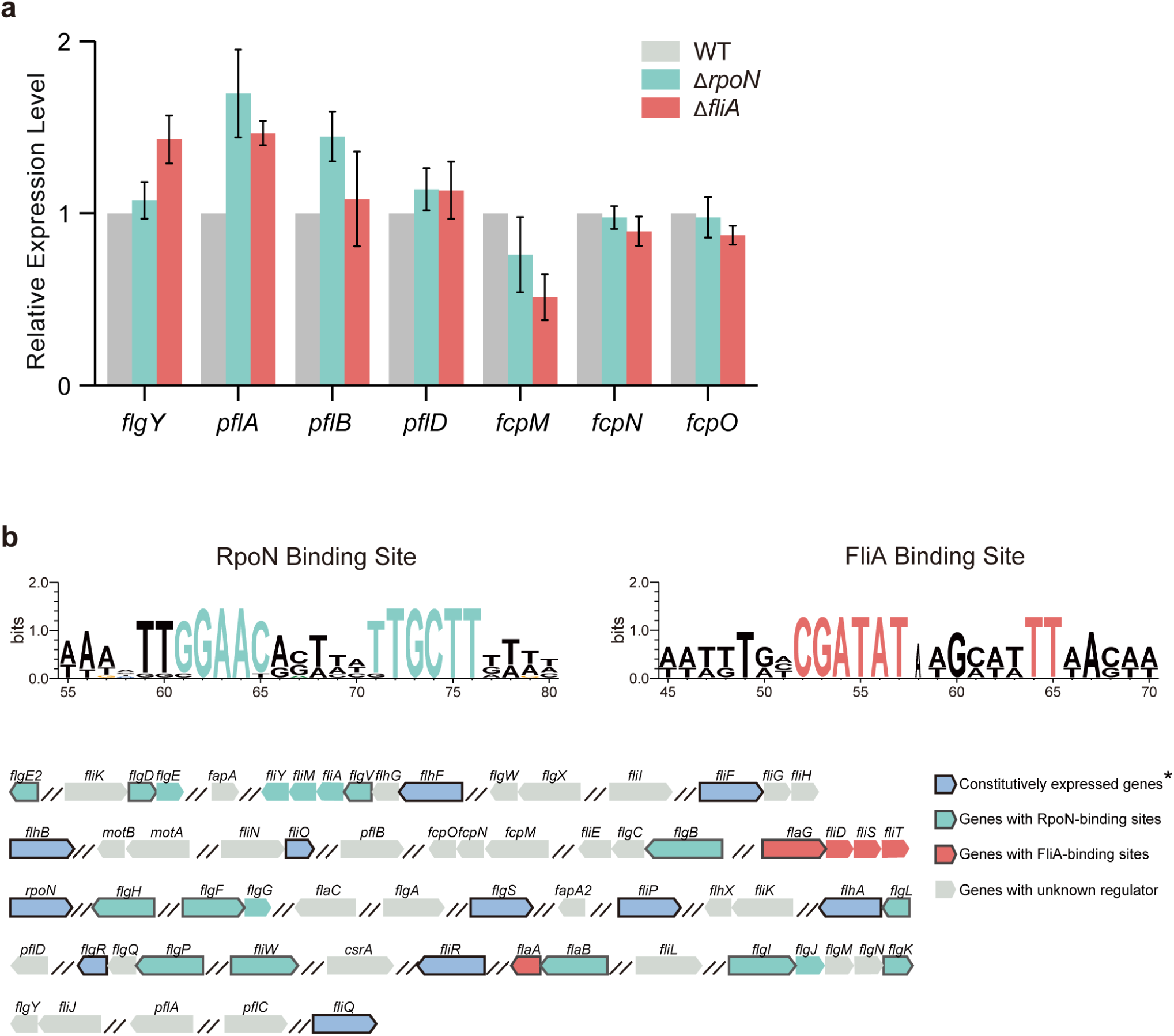
The regulation of *flgY*, *pflA*, *pflB*, *pflD*, and *fcpMNO*. **a**, qPCR analyses of expression level of seven genes from Δ*rpoN* or Δ*fliA* mutant compared to WT. Data are presented as mean from three independent experiments. No statistically significant differences were observed compared to the wild-type control at the threshold of p-value < 0.001. **b**, Promoter region analyses of flagellar genes/operons in *C. jejuni* 81-176 to show the presence or absence of RpoN- or FliA- binding sites. The RpoN- or FliA- binding motif shown on the top are taken from reference ^1^. Constitutively expressed genes marked with an asterisk are information adopted from reference ^2,3^.

**Extended Data Figure 9.**
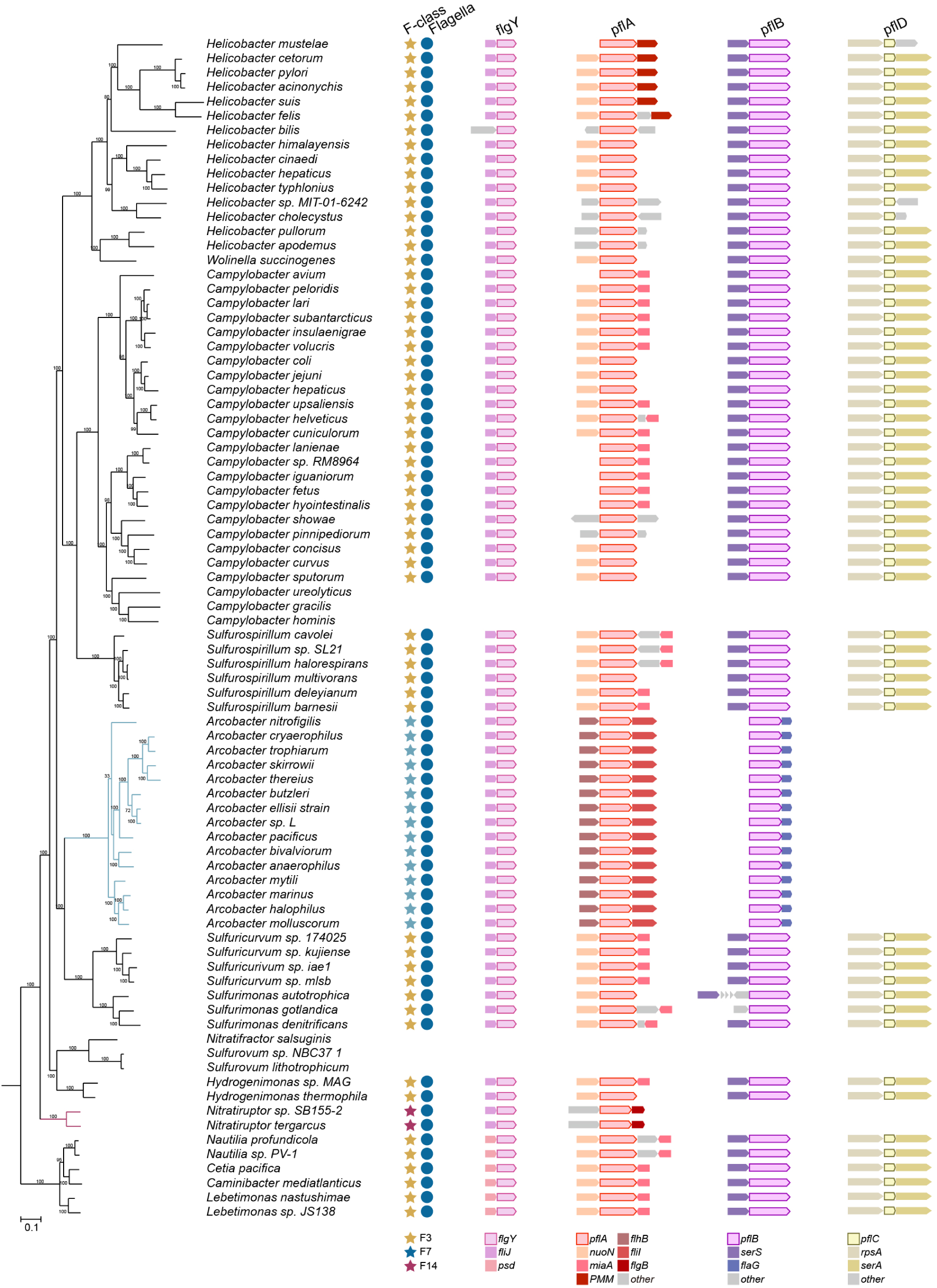
The conserved gene order of *flgY*, *pflA*, *pflB*, and *pflD* in *Campylobacterota* genomes. Branches were highlighted for *Arcobacter* (blue) and *Nitratiruptor* (red) that have non-F3 chemosensory class.

**Extended Data Figure 10.**
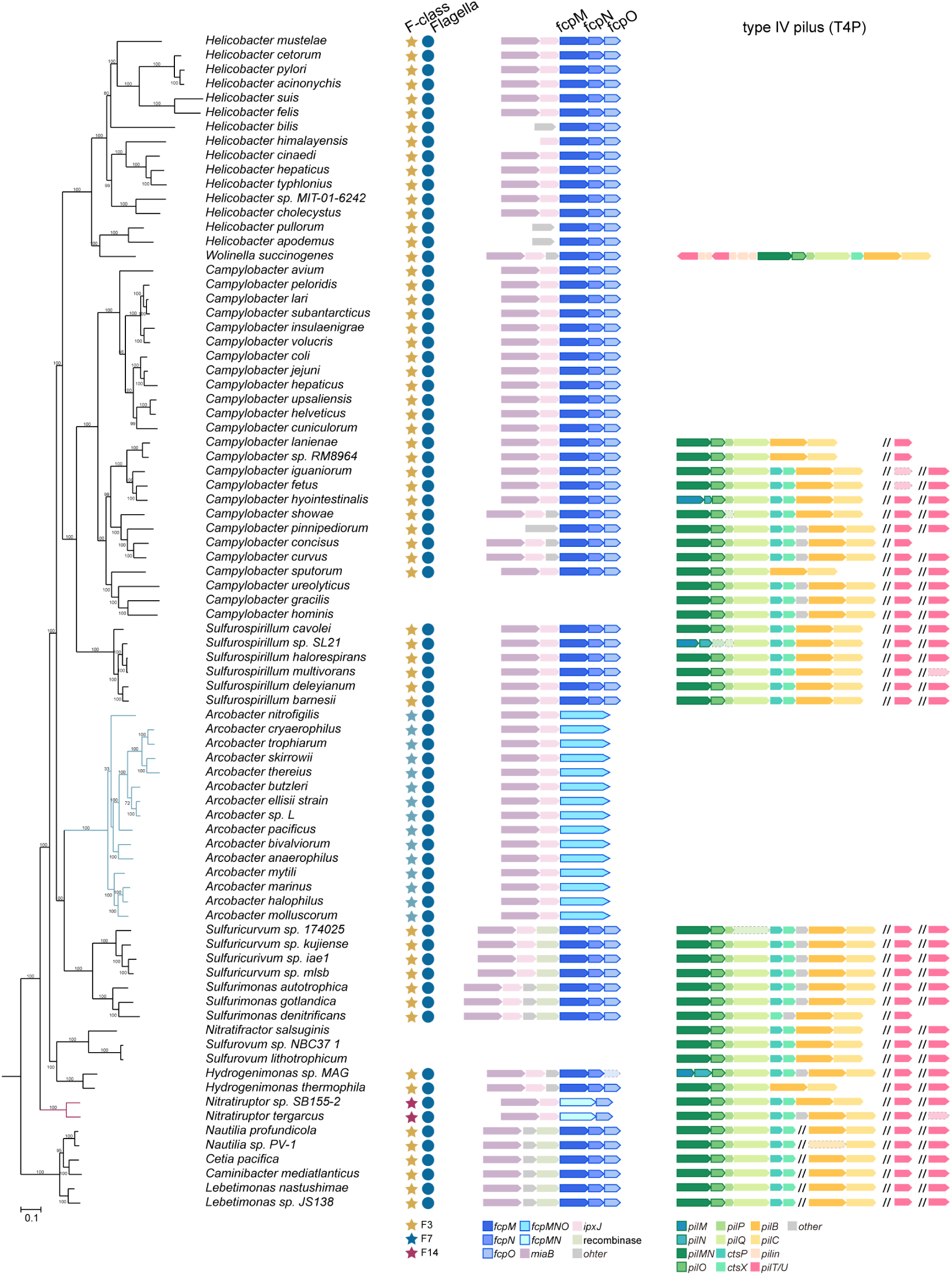
The conserved gene order of *fcpMNO operon* with its upstream genes in *Campylobacterota* genomes. The identified gene clusters of T4P in *Campylobacterota* genomes are also shown here. Branches were highlighted for *Arcobacter* (blue) and *Nitratiruptor* (red) that have non-F3 chemosensory class and their genes of *fcpMNO* operon have fusion as indicated at the bottom of the figure.

**Extended Data Figure 11.**
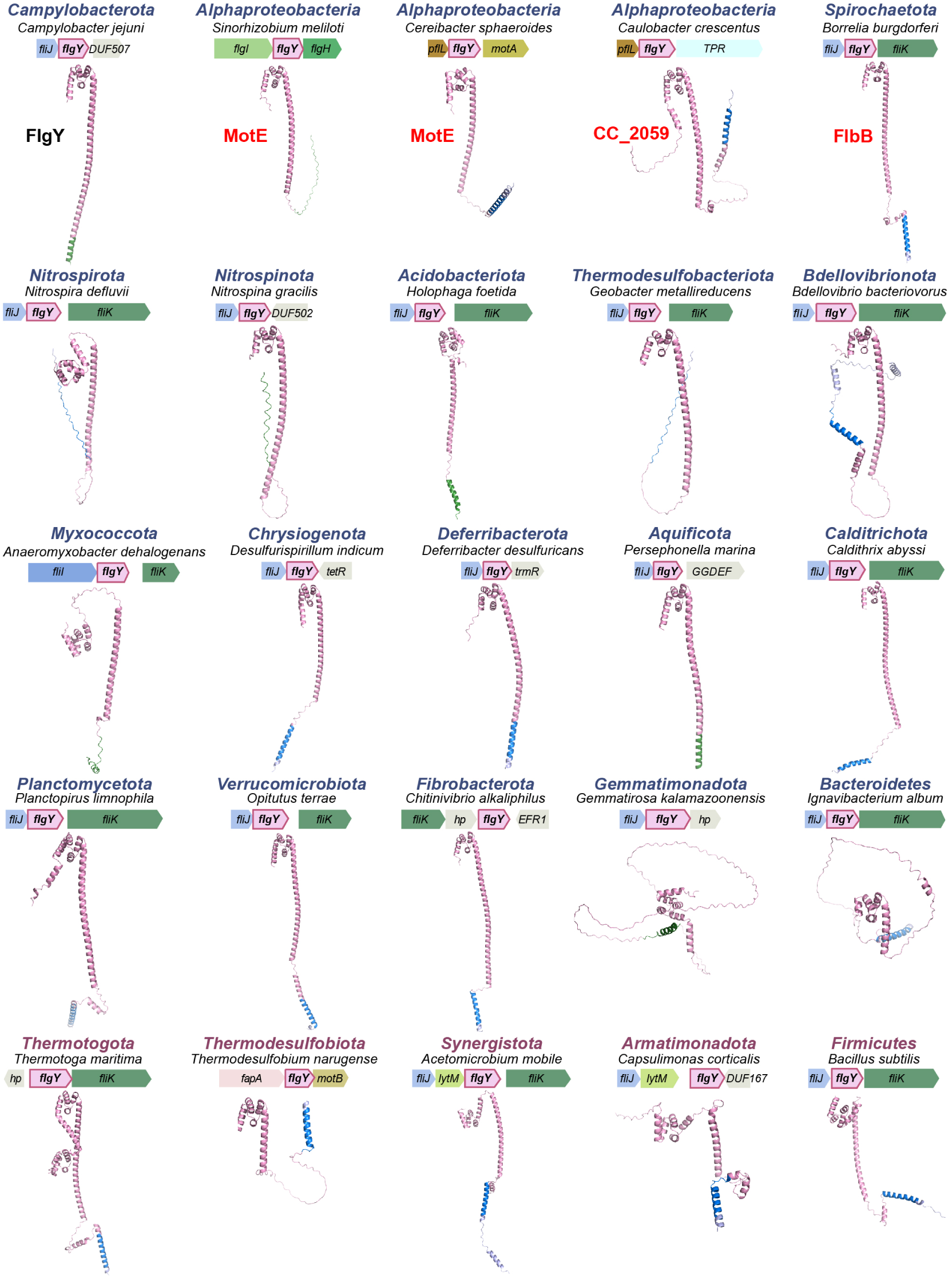
Structures of FlgY homologs and associated genomic contexts for representative species across phyla. The transmembrane motif and signal peptide predicted by TMHMM 2.0 and SignalP 6.0 are colored in blue and green, respectively. All structures are predicted by AlphaFold3. The protein names highlighted in red on the top row are FlgY homologs that have been experimentally characterized, including MotE from both *S. meliloti* ^4^ and *C. sphaeroides* ^5^ and FlbB from *B. burgdorferi* ^6^. CC_2059 from *C. crescentus* is also highlighted in red since the E-ring is first identified from this species ^7,8^ and most likely CC_2059 constitutes the E-ring.

**Extended Data Fig Figure 12.**
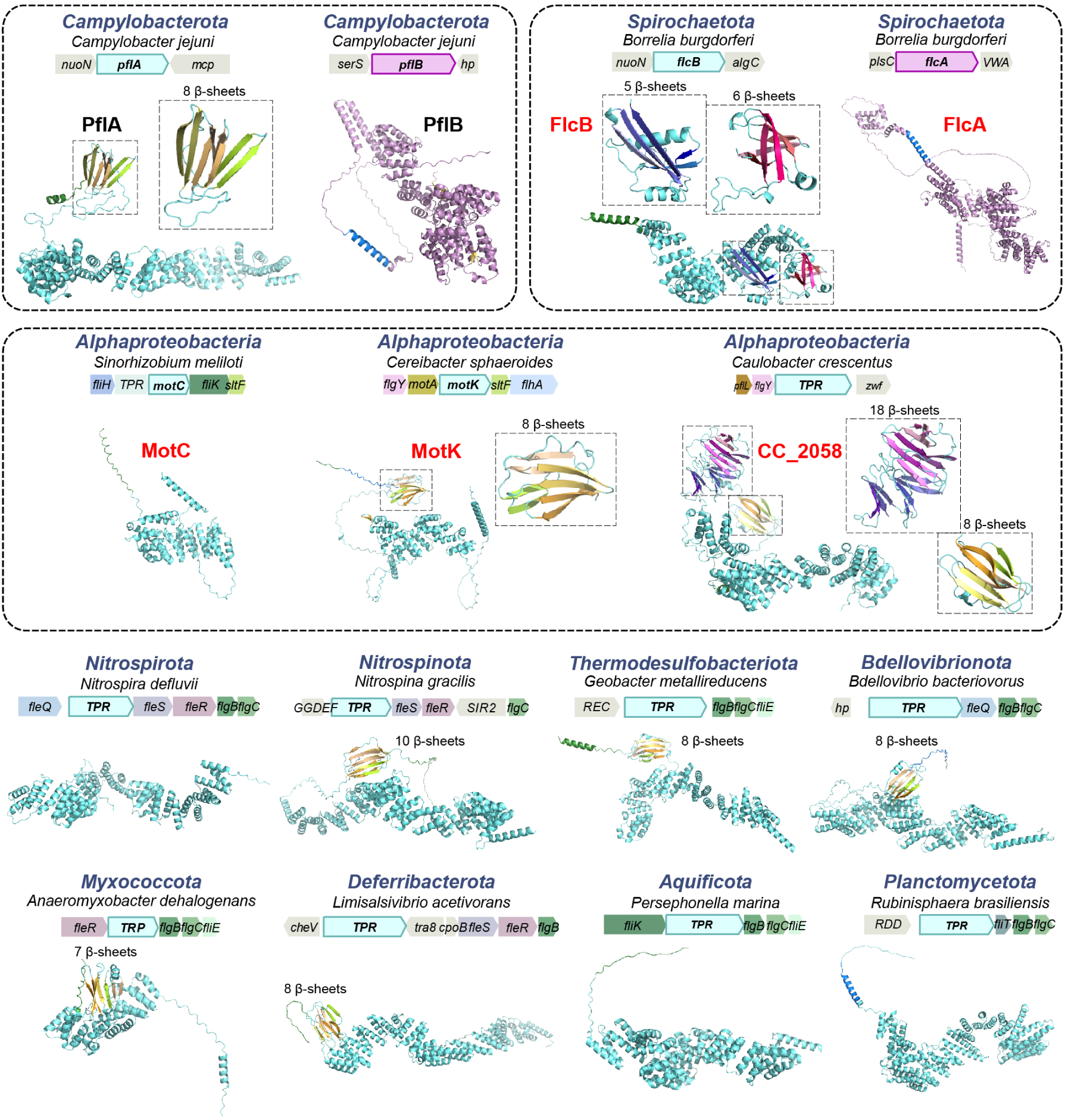
Structures of PflAB homologs and associated genomic contexts for representative species across phyla. The transmembrane motif and signal peptide predicted by TMHMM 2.0 and SignalP 6.0 are colored in blue and green, respectively. The number of β-sheets in the β-sandwich domain is indicated above each structure. All structures are predicted by AlphaFold3. The protein names highlighted in red on the top rows are PflA or PflB homologs that have been experimentally characterized, including MotC from *S. meliloti* ^4^, MotK from *C. sphaeroides* ^5^ and FlcAB from *B. burgdorferi* ^9,10^. CC_2058 from *C. crescentus* is also highlighted in red since most likely CC_2058 interacts with CC_2059 that constitutes the E- ring.

**Extended Data Figure 13.**
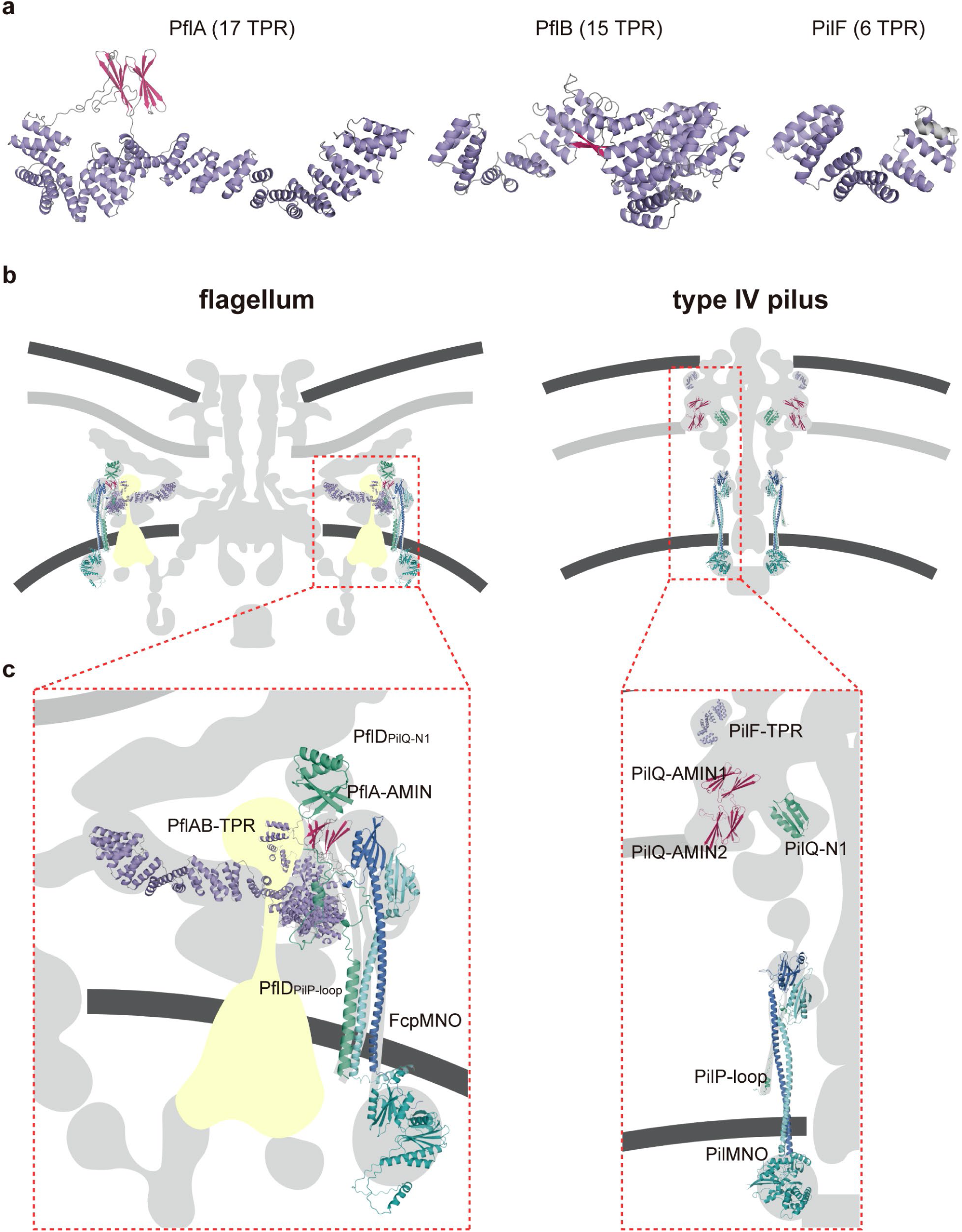
Structural Comparisons of homologous proteins in flagellum and T4P. **a,** Structural comparison of PflA, PflB, and PilF (PDB:2FI7). **b-c**, Scaffolding components of *C. jejuni* motor share homology with T4P proteins.

**Extended Data Fig Figure 14.**
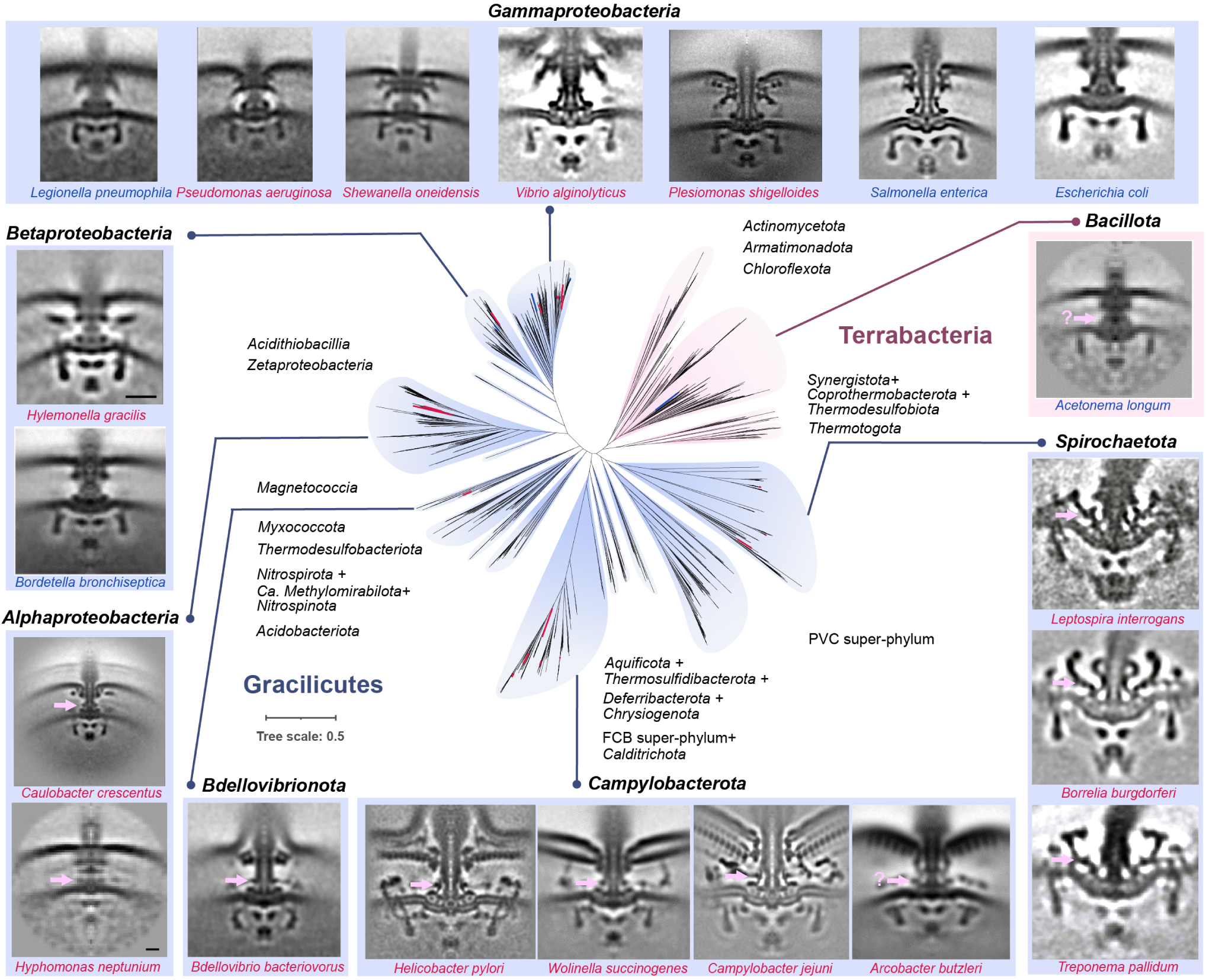
Summary of motor structures imaged by cryo-ET and mapped to bacterial species tree. The phylogenetic tree in the center is derived from Fig. 6c. The motor structures investigated by cryo-ET are displayed on the periphery, with one representative species from each genus and taxon group labeled at the top of the image. Species names highlighted in blue indicate that their stator complexes are dynamic and invisible, while those marked in red represent that their stator complexes are visible. Potential E- ring is indicated by pink arrow and question marks in the image of *Acetonema longum* and *Arcobacter butzleri* mean that the position of E-ring is uncertain. Information of motor structure for species (follow a clockwise order) were taken from references: *Acetonema longum* ^11^, *Leptospira interrogans* ^12^, *Borrelia burgdorferi* ^13^, *Treponema pallidum* ^14^, *Arcobacter butzleri* ^15^, *Campylobacter jejuni* (this study), *Wolinella succinogenes* ^15^, *Helicobacter pylori* ^16^, *Bdellovibrio bacteriovorus* ^15^, *Hyphomonas neptunium* ^11^, *Caulobacter crescentus* ^17^, *Bordetella bronchiseptica* ^18^, *Hylemonella gracilis* ^11^, *Legionella pneumophila* ^19^, *Pseudomonas aeruginosa* ^19^, *Shewanella oneidensis* ^19^, *Vibrio alginolyticus* ^20^, *Plesiomonas shigelloides* ^21^, *Salmonella enterica* ^22^, *Escherichia coli* ^23^.

